# Molecular dissection of PI3Kβ synergistic activation by receptor tyrosine kinases, GβGγ, and Rho-family GTPases

**DOI:** 10.1101/2023.05.01.538969

**Authors:** Benjamin R. Duewell, Naomi E. Wilson, Gabriela M. Bailey, Sarah E. Peabody, Scott D. Hansen

## Abstract

The class 1A phosphoinositide 3-kinase (PI3K) beta (PI3Kβ) is functionally unique in the ability to integrate signals derived from receptor tyrosine kinases (RTKs), heterotrimeric guanine nucleotide-binding protein (G-protein)-coupled receptors (GPCRs), and Rho-family GTPases. The mechanism by which PI3Kβ prioritizes interactions with various membrane tethered signaling inputs, however, remains unclear. Previous experiments have not been able to elucidate whether interactions with membrane-tethered proteins primarily control PI3Kβ localization versus directly modulate lipid kinase activity. To address this gap in our understanding of PI3Kβ regulation, we established an assay to directly visualize and decipher how three distinct protein interactions regulate PI3Kβ when presented to the kinase in a biologically relevant configuration on supported lipid bilayers. Using single molecule Total Internal Reflection Fluorescence (TIRF) Microscopy, we determined the mechanism controlling membrane localization of PI3Kβ, prioritization of signaling inputs, and lipid kinase activation. We find that auto-inhibited PI3Kβ prioritizes interactions with RTK-derived tyrosine phosphorylated (pY) peptides before engaging either GβGγ or Rac1(GTP). Although pY peptides strongly localize PI3Kβ to membranes, stimulation of lipid kinase activity is modest. In the presence of either pY/GβGγ or pY/Rac1(GTP), PI3Kβ activity is dramatically enhanced beyond what can be explained by simply increasing the strength of membrane localization. Instead, PI3Kβ is synergistically activated by pY/GβGγ and pY/Rac1(GTP) through a mechanism consistent with allosteric regulation.

## INTRODUCTION

Critical for cellular organization, phosphatidylinositol phosphate (PIP) lipids regulate the localization and activity of numerous proteins across intracellular membranes in eukaryotic cells (Di Paolo and De Camilli 2006). The interconversion between various PIP lipid species through the phosphorylation and dephosphorylation of inositol head groups is regulated by lipid kinases and phosphatases (Balla 2013; Burke 2018). Serving a critical role in cell signaling, the class I family of phosphoinositide 3-kinases (PI3Ks) catalyze the phosphorylation of phosphatidylinositol 4,5-bisphosphate [PI(4,5)P_2_] to generate PI(3,4,5) P_3_. Although a low-abundance lipid (< 0.05%) in the plasma membrane (Wenk et al. 2003; Nasuhoglu et al. 2002; Stephens, Jackson, and Hawkins 1993), PI(3,4,5)P_3_ can increase 40-fold following receptor activation (Stephens, Hughes, and Irvine 1991; Parent et al. 1998; Insall and Weiner 2001). Although signal adaptation mechanisms typically restore PI(3,4,5)P_3_ to the basal level following receptor activation (Funamoto et al. 2002; Yip et al. 2008; Auger et al. 1989), misregulation of the PI3K signaling pathway can result in constitutively high levels of PI(3,4,5)P_3_ that are detrimental to cell health. Since PI(3,4,5)P_3_ lipids serve an instructive role in driving actin based membrane protrusions (Howard and Oresajo 1985; Weiner 2002; Graziano et al. 2017), sustained PI(3,4,5)P_3_ signaling is known to drive cancer cell metastasis (Hanker et al. 2013). Elevated PI(3,4,5)P_3_ levels also stimulates the AKT signaling pathway and Tec family kinases, which can drive cellular proliferation and tumorigenesis (Manning and Cantley 2007; Fruman et al. 2017). While much work has been dedicated in determining the factors that participate in the PI3K signaling pathway, how these molecules collaborate to rapidly synthesize PI(3,4,5)P_3_ remains an important open question. To decipher how amplification of PI(3,4,5)P_3_ arises from the relay of signals between cell surface receptors, lipids, and peripheral membrane proteins, we must understand how membrane localization and activity of PI3Ks is regulated by different signaling inputs. Determining how these biochemical reactions are orchestrated will provide new insight concerning the molecular basis of asymmetric cell division, cell migration, and tissue organization, which are critical for understanding development and tumorigenesis.

In the absence of a stimulatory input, the class IA family of PI3Ks (PI3Kα, PI3Kβ, PI3Kδ) are thought to reside in the cytoplasm as auto-inhibited heterodimeric protein complexes composed of a catalytic (p110α, p110β, or p110δ) and regulatory subunit (p85α, p85β, p55γ, p50α, or p55α) (Burke 2018; Vadas et al. 2011). The catalytic subunits of class IA PI3Ks contain an N-terminal adaptor binding domain (ABD), a Ras/Rho binding domain (RBD), a C2 domain (C2), and an adenosine triphosphate (ATP) binding pocket (Vadas et al. 2011). The inter-SH2 (iSH2) domain of the regulatory subunit tightly associates with the ABD of the catalytic subunit (Yu et al. 1998), providing structural integrity, while limiting dynamic conformational changes. The nSH2 and cSH2 domains of the regulatory subunit form additional inhibitory contacts that limit the conformational dynamics of the catalytic subunit (Zhang et al. 2011; Mandelker et al. 2009; Burke et al. 2011; Carpenter et al. 1993; Yu et al. 1998). A clearer understanding of the how various proteins control PI3K localization and activity would help facilitate the development of drugs that perturb specific protein-protein binding interfaces that are critical for membrane targeting and lipid kinase activity.

Among the class IA PI3Ks, PI3Kβ is uniquely capable of interacting with Rho-family GTPases (Fritsch et al. 2013), Rab GTPases (Christoforidis et al. 1999; Heitz et al. 2019), heterotrimeric G-protein complexes (GβGγ) (Kurosu et al. 1997; Maier, Babich, and Nürnberg 1999; Guillermet-Guibert et al. 2008), and phosphorylated receptor tyrosine kinases (RTKs) (Zhang et al. 2011; Carpenter et al. 1993). Like other class IAPI3Ks, interactions with receptor tyrosine kinase (RTK) derived phosphotyrosine peptides release nSH2 and cSH2-mediated inhibition of the catalytic subunit to stimulate PI3Kβ lipid kinase activity (H. A. Dbouk et al. 2012; Zhang et al. 2011). GβGγ and Rac1(GTP) in solution have also been shown to stimulate PI3Kβ lipid kinase activity (Hashem A. Dbouk et al. 2012; Fritsch et al. 2013; Maier, Babich, and Nürnberg 1999). Similarly, activation of Rho-family GTPases (Fritsch et al. 2013) and G-protein coupled receptors (Houslay et al. 2016) stimulate PI3Kβ lipid kinase activity in cells. However, it’s unclear how individual interactions with GβGγ or Rac1(GTP) can bypass autoinhibition of full-length PI3Kβ (p110β-p85α/β). Studies in neutrophils and *in vitro* biochemistry suggest that PI3Kβ is synergistically activated through coincidence detection of RTKs and GβGγ (Houslay et al. 2016; Hashem A. Dbouk et al. 2012). Similarly, Rac1(GTP) and GβGγ have been reported to synergistically activate PI3Kβ in cells (Erami et al. 2017). An enhanced membrane recruitment mechanism is the most prominent model used to explain synergistic activation of PI3Ks.

There is limited kinetic data examining how PI3Kβ is regulated by different membrane-tethered proteins. Previous biochemical studies of PI3Kβ have utilized solution-based assays to measure P(3,4,5)P_3_ production. As a result, the mechanisms that determine how PI3Kβ prioritizes interactions with RTKs, small GTPases, or GβGγ remains unclear. In the case of synergistic PI3Kβ activation, it’s unclear which protein-protein interactions regulate membrane localization versus stimulate lipid kinase activity. No studies have simultaneously measured PI3Kβ membrane association and lipid kinase activity to decipher potential mechanisms of allosteric regulation. Previous studies concerning the synergistic activation of PI3Ks are challenging to interpret because RTK derived peptides are always presented in solution alongside membrane anchored signaling inputs. However, all the common signaling inputs for PI3K activation (i.e. RTKs, GβGγ, Rac1/Cdc42) are membrane associated proteins. Activation of class 1A PI3Ks has never been reconstituted using solely membrane tethered activators conjugated to membranes in a biologically relevant configuration. As a result, we currently lack a comprehensive description of PI3Kβ membrane recruitment and catalysis.

To decipher the mechanisms controlling PI3Kβ membrane binding and activation, we established a biochemical reconstitution using supported lipid bilayers (SLBs). We used single molecule Total Internal Reflection Fluorescence (TIRF) microscopy to quantify the relationship between PI3Kβ localization, lipid kinase activity, and the density of various membrane-tethered signaling inputs. This approach allowed us to measure the dwell time, binding frequency, and diffusion coefficients of single fluorescently labeled PI3Kβ in the presence of RTK derived peptides, Rac1(GTP), and GβGγ. Simultaneous measurements of PI3Kβ membrane recruitment and lipid kinase activity allowed us to define the relationship between PI3Kβ localization and PI(3,4,5)P_3_ production in the presence of different regulators. Overall, we found that membrane docking of PI3Kβ first requires interactions with RTK-derived tyrosine phosphorylated (pY) peptides, while PI3Kβ localization is insensitive to membranes that contain either Rac1(GTP) or GβGγ alone. Following engagement with a pY peptide, PI3Kβ can associate with either GβGγ or Rac1(GTP). In the case of synergistic PI3Kβ localization mediated by pY/GβGγ, it’s essential for the nSH2 domain to move away from the GβGγ binding site. Although both the PI3Kβ-pY-Rac1(GTP) and PI3Kβ-pY-GβGγ complexes display a ∼2-fold increase in membrane localization, the corresponding increase in catalytic efficiency is much greater. Overall, our results indicate that synergistic activation of PI3Kβ depends on allosteric modulation of lipid kinase activity.

## RESULTS

### PI3Kβ prioritizes interactions with pY peptides over Rac1(GTP) and GβGγ

Previous biochemical analysis of p110β-p85α, referred to as PI3Kβ, established that receptor tyrosine kinases (Zhang et al. 2011), Rho-type GTPases (Fritsch et al. 2013), and heterotrimeric G-protein GβGγ (Hashem A. Dbouk et al. 2012) are capable of binding and stimulating lipid kinase activity. To decipher how PI3Kβ prioritizes interactions between these three membrane-tethered proteins we established a method to directly visualize PI3Kβ localization on supported lipid bilayers (SLBs) using Total Internal Reflection Fluorescence (TIRF) Microscopy (**Figure 1A**). For this assay, we covalently attached either a doubly tyrosine phosphorylated platelet derived growth factor (PDGF) peptide (pY) peptide or recombinantly purified Rac1 to supported membranes using cysteine reactive maleimide lipids. We confirmed membrane conjugation of the pY peptide and Rac1 by visualizing the localization of fluorescently labeled nSH2-Cy5 or Cy3-p67/phox (Rac1(GTP) sensor), respectively (**Figure 1B**). Nucleotide exchange of membrane conjugated Rac1(GDP) was achieved by the addition of a guanine nucleotide exchange factor, P-Rex1 (phosphatidylinositol 3,4,5-trisphosphate-dependent Rac exchanger 1 protein) diluted in GTP containing buffer (**Figure 1C**). As previously described (Rathinaswamy et al. 2021; 2023), AF488-SNAP dye labeled farnesyl GβGγ was directly visualized following passive absorption into supported membranes (**Figure 1D** and **Figure 1 – figure supplement 1**). We confirmed that membrane bound GβGγ was functional by visualizing robust membrane recruitment of Dy647-PI3Kγ by TIRF-M (**Figure 1E**). Overall, this assay functions as a mimetic to the cellular plasma membrane and allowed us to examine how different membrane tethered signaling inputs regulate PI3Kβ membrane localization in vitro.

**Figure 1.**
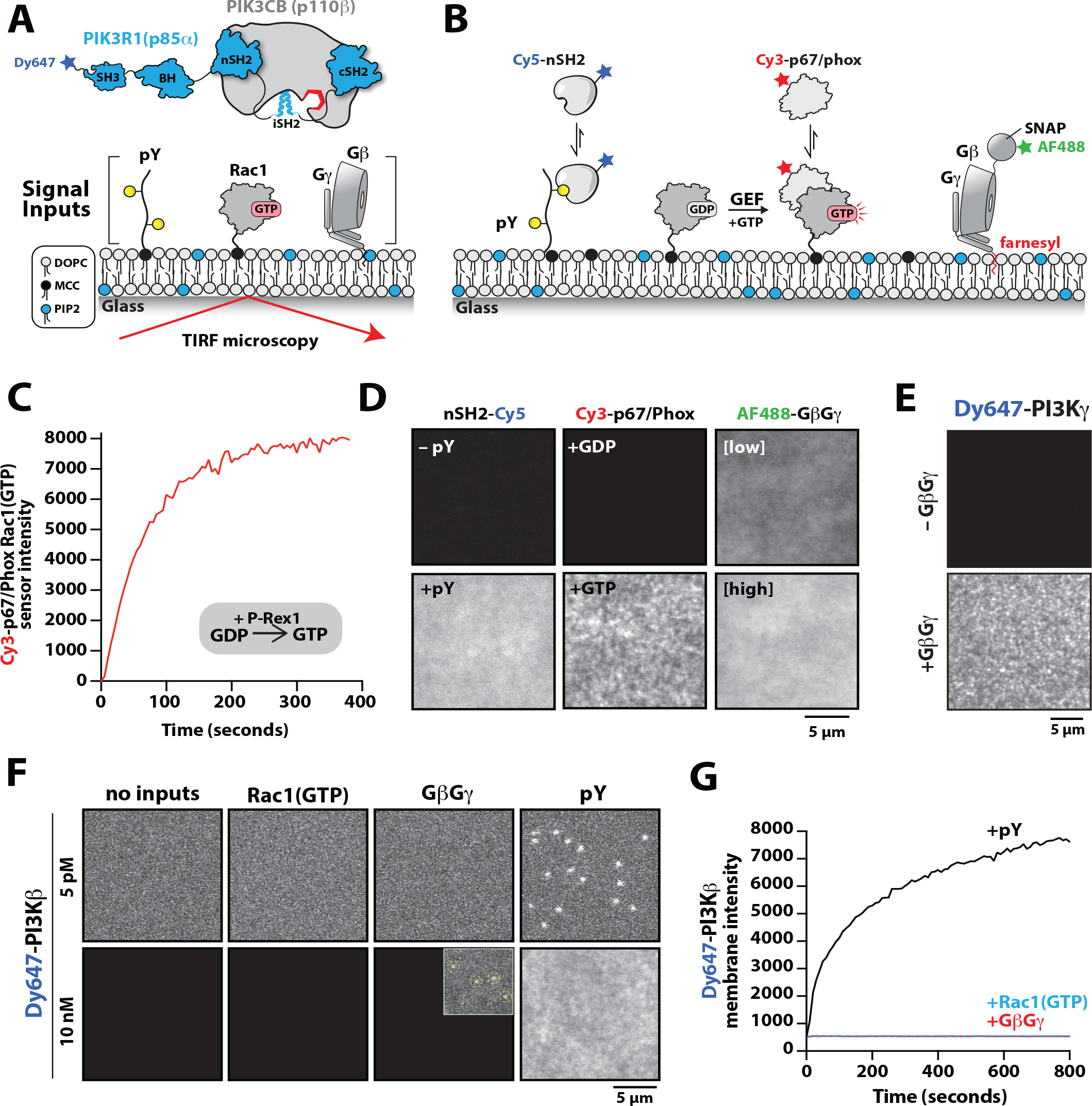
PI3Kβ prioritizes membrane interactions with RTK-derived pY peptides over Rac1(GTP) and GβGγ. **(A)** Cartoon schematic showing membrane tethered signaling inputs (i.e. pY, Rac1(GTP), and GβGγ) attached to a supported lipid bilayer and visualized by TIRF-M. Heterodimeric Dy647-PI3Kβ (p110β-p85α) in solution can dynamically associate with membrane bound proteins. **(B)** Cartoon schematic showing method for visualizing membrane tethered signaling inputs. **(C)** Kinetics of Rac1 nucleotide exchange measured in the presence of 20 nM Rac1(GTP) sensor (Cy3-p67/phox) and 50 nM P-Rex1. **(D)** Visualization of membrane conjugated RTK derived pY peptide (∼6,000/μm^2^), Rac1(GTP) (∼4,000/μm^2^), and GβGγ (∼4,800/μm^2^) by TIRF-M. Representative TIRF-M images showing the membrane localization of 20 nM nSH2-Cy3 in the absence and presence of membranes conjugated with a solution concentration of 10 μM pY peptide. Representative images showing the membrane localization of 20 nM Cy3-p67/phox Rac1(GTP) sensor before (GDP) and after (GTP) the addition of the guanine nucleotide exchange factor, P-Rex1. Equilibrium localization of 50 nM (low) or 200 nM (high) farnesyl GβGγ-SNAP-AF488. **(E)** Representative TIRF-M images showing the equilibrium membrane localization of 10 nM Dy647-PI3Kγ measured in the absence and presence of membranes equilibrated with 200 nM farnesyl GβGγ. **(F)** Representative TIRF-M images showing the equilibrium membrane localization of 5 pM and 10 nM Dy647-PI3Kβ measured in the presence of membranes containing either pY, Rac1(GTP), or GβGγ. The inset image (+GβGγ) shows low frequency single molecule binding events detected in the presence of 10 nM Dy647-PI3Kβ. **(G)** Bulk membrane absorption kinetics for 10 nM Dy647-PI3Kβ measured on membranes containing either pY, Rac1(GTP), or GβGγ. Membrane composition: 96% DOPC, 2% PI(4,5)P_2_, 2% MCC-PE.

We visualized both single molecule binding events and bulk membrane localization of Dy647-PI3Kβ by TIRF-M to determine which inputs can autonomously recruit autoinhibited Dy647-PI3Kβ from solution to a supported membrane (**Figure 1F**). Comparing membrane localization of Dy647-PI3Kβ in the presence of pY, Rac1(GTP), or GβGγ revealed that only the tyrosine phosphorylated peptide (pY) could robustly localize Dy647-PI3Kβ to supported membranes (**Figure 1F-1G**). This prioritization of interactions was consistently observed across a variety of membrane lipid compositions (**Figure 1 – figure supplement 2**). Increasing the anionic lipid charge of supported membranes through the addition of 20% phosphatidylserine (PS), did not significantly change the frequency Dy647-PI3Kβ membrane interactions in the presence of either Rac1(GTP) or GβGγ alone (**Figure 1 – figure supplement 2**). Although we could detect some transient Dy647-PI3Kβ membrane binding events in the presence of GβGγ alone, the binding frequency was reduced 2000-fold compared our measurements on pY membranes (**Figure 1 – figure supplement 2**). Localization of wild-type Dy647-PI3Kβ phenocopied the GβGγ binding mutant, Dy647-PI3Kβ (K532D, K533D), indicating that the low frequency binding events we observed are mostly mediated by lipid interactions rather than direct binding to GβGγ (**Figure 1 – figure supplement 2**). The addition of 20% PS lipids did, however, cause a subtle increase in the number of Dy647-PI3Kβ membrane binding observed when 20 μM pY peptide was added in solution (**Figure 1 – figure supplement 2**). Our observations are consistent with previous reports involving the membrane binding dynamics of AF555-PI3Kα measured in the presence of solution pY peptide (Ziemba et al. 2016; Buckles et al. 2017). Given the large parameter space, we did not perform an exhaustive characterization of Dy647-PI3Kβ membrane binding across many membrane compositions. In this study, we use a simplified membrane composition that minimized non-specific membrane localization of fluorescently labeled PI3Kβ. This allowed us to more clearly define the strength of individual and combinations of protein-protein interactions that regulate PI3Kβ localization and kinase activity.

### PI3Kβ cooperatively engages a single membrane tethered pY peptide

Previous biochemical analysis of PI3Kβ utilized pY peptides in solution to study the regulation of lipid kinase activity (Zhang et al. 2011; Hashem A. Dbouk et al. 2012). Using membrane-tethered pY peptide, we quantitatively mapped the relationship between the pY membrane surface density and the membrane binding behavior of Dy647-PI3Kβ (**Figure 2A**). To calculate the membrane surface density of conjugated pY, we incorporated a defined concentration of Alexa488-pY (**Figure 2A-2B**). We measured the relationship between the total solution concentration of pY peptide used for the membrane conjugation step and the corresponding final membrane surface density (pY per μm^2^). Over a range of pY peptide solution concentrations (0-10 μM), we observed a linear increase in the membrane conjugation efficiency based on the incorporation of fluorescent Alexa488-pY (**Figure 2C, left hand y-axis**). Bulk membrane localization of a nSH2-Cy3 sensor showed a corresponding linear increase in fluorescence as a function of pY peptide membrane density (**Figure 2D**). By quantifying the average number of Alexa488-pY particles per unit area of supported membrane we calculated the absolute density of pY per μm^2^ (**Figure 2C, right hand y-axis**).

**Figure 2.**
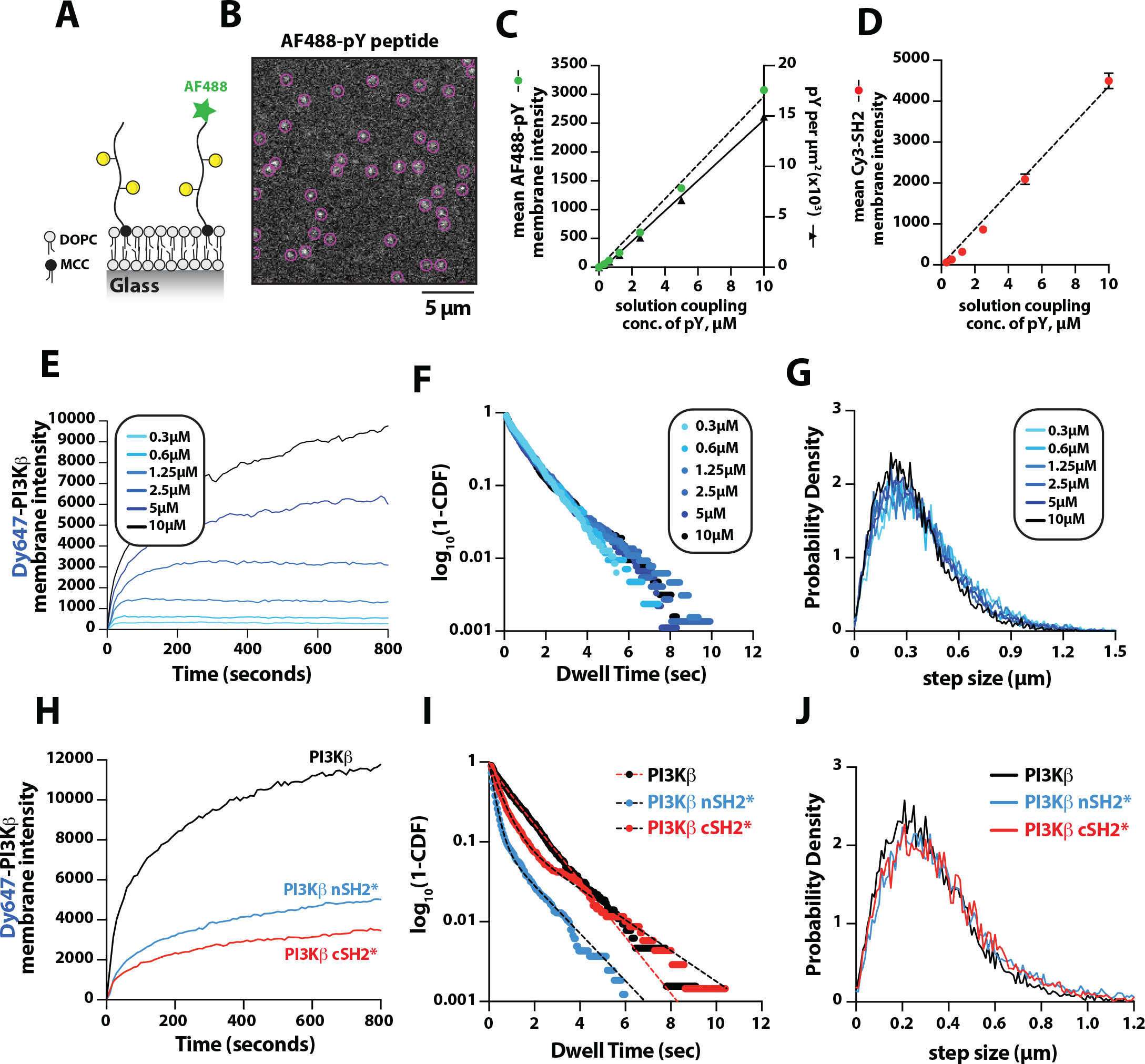
Density dependent membrane binding behavior of Dy647-PI3Kβ measured in the presence of RTK-derived pY peptides. **(A)** Cartoon schematic showing conjugation of pY peptides (+/-Alexa488 label) using thiol reactive maleimide lipids (MCC-PE). **(B)** Representative image showing the single molecule localization of Alexa488-pY. Particle detection (purple circles) was used to quantify the number of pY peptides per μm^2^. **(C)** Relationship between the total pY solution concentration (x-axis) used for covalent conjugation, the bulk membrane intensity of covalently attached Alexa488-pY (left y-axis), and the final surface density of pY peptides per μm^2^ (right y-axis). **(D)** Relationship between the total pY solution conjugation concentration and bulk membrane intensity of measured in the presence of 50 nM nSH2-Cy3. **(E-G)** Membrane localization dynamics of Dy647-PI3Kβ measured on SLBs containing a range of pY surface densities (250–15,000 pY/μm^2^, based on Figure 1C). **(E)** Bulk membrane localization of 10 nM Dy647-PI3Kβ as a function of pY density. **(F)** Single molecule dwell time distributions measured in the presence of 5 pM Dy647-PI3Kβ. Data plotted as log_10_(1–CDF) (cumulative distribution frequency). **(G)** Step size distributions showing Dy647-PI3Kβ single molecule displacements from > 500 particles (>10,000 steps) per pY surface density. **(H-J)** Membrane localization dynamics of Dy647-PI3Kβ nSH2(R358A) and cSH2(R649A) mutants measured on SLBs containing ∼15,000 pY/μm^2^ (10μM conjugation concentration). **(H)** Bulk membrane absorption kinetics of 10 nM Dy647-PI3Kβ (WT, nSH2*, and cSH2*). **(I)** Single molecule dwell time distributions measured in the presence of 5 pM Dy647-PI3Kβ (WT, nSH2*, and cSH2*). Data plotted as log_10_(1–CDF) (cumulative distribution frequency). **(J)** Step size distributions showing single molecule displacements of > 500 particles (>10,000 steps) in the presence of 5 pM Dy647-PI3Kβ (WT, nSH2*, and cSH2*). Membrane composition: 96% DOPC, 2% PI(4,5)P_2_, 2% MCC-PE.

To determine how the membrane binding behavior of PI3Kβ is modulated by the membrane surface density of pY, we measured the bulk membrane absorption kinetics of Dy647-PI3Kβ. When Dy647-PI3Kβ was flowed over membranes containing a surface density between 250 and 3,000 pY/μm^2^, we observed rapid equilibration kinetics consistent with a simple biomolecular interaction (**Figure 2E**). Similar membrane binding kinetics have been reported for the Btk-PI(3,4,5)P_3_ (Chung et al. 2019) and PI3Kγ-GβGγ (Rathinaswamy et al. 2021) complexes. When the membrane surface density was greater than 6,500 pY/μm^2^, we observed slower equilibration kinetics consistent with a fraction of Dy647-PI3Kβ complexes exhibiting longer dwell times. Single particle tracking of Dy647-PI3Kβ on membranes containing varying densities of pY peptide revealed that the dwell time was relatively insensitive to the pY peptide density (**Figure 2F** and **Table 1**). However, we could be underestimating the actual dwell time of Dy647-PI3Kβ due to membrane hopping (Yasui, Matsuoka, and Ueda 2014). In the presence of a high pY surface density, Dy647-PI3Kβ could dissociate from pY and then immediately rebind to another pY instead of diffusing into the solution phase. Quantification of single particle displacement (or step size) of pY-tethered Dy647-PI3Kβ complexes revealed nearly identical diffusivity across a range of pY membrane densities (**Figure 2G** and **Table 1**). Together, these results suggest that Dy647-PI3Kβ most frequently engages a single dually phosphorylated peptide over a broad range of pY densities in our bilayer assay.

The regulatory subunit of PI3Kβ (p85α) contains two SH2 domains that form inhibitory contacts with the catalytic domain (p110β) (Zhang et al. 2011). The SH2 domains of class 1A PI3Ks have a conserved peptide motif, FLVR, that mediates the interaction with tyrosine phosphorylated peptides (Bradshaw, Mitaxov, and Waksman 1999; Waksman et al. 1992; Rameh, Chen, and Cantley 1995). Mutating the arginine to alanine (FLVA mutant) prevents the interaction with pY peptides for both PI3Kα and PI3Kβ (Yu et al. 1998; Dornan et al. 2020; Nolte et al. 1996; Zhang et al. 2011; Breeze et al. 1996). To determine how the membrane binding behavior of PI3Kβ is modulated by each SH2 domain, we individually mutated the FLVR amino acid sequence to FLVA. Compared to wild type Dy647-PI3Kβ, the nSH2(R358A) and cSH2(R649A) mutants showed a 60% and 75% reduction in membrane localization at equilibrium, respectively (**Figure 2H**). Single molecule dwell time analysis also showed a significant reduction in membrane affinity for Dy647-PI3Kβ nSH2(R358A) and cSH2(R649A) compared to wild type PI3Kβ (**Figure 2I** and **Table 1**). Single molecule diffusion (or mobility) of membrane bound nSH2(R358A) and cSH2(R649A) mutants, however, were nearly identical to wild type Dy647-PI3Kβ (**Figure 2J** and **Table 1**). Because the nSH2 and cSH2 mutants can only interact with a single phosphorylated tyrosine residue on the doubly phosphorylated pY peptide, this further supports a model in which the p85α regulatory subunit of PI3Kβ cooperatively engages one doubly phosphorylated pY peptide under our experimental conditions.

### GβGγ dependent enhancement in PI3Kβ localization requires release of the nSH2

Having established that PI3Kβ engagement with a membrane tethered pY peptide is the critical first step for robust membrane localization, we examined the secondary role that GβGγ serves in controlling membrane localization of PI3Kβ bound to pY. To measure synergistic membrane localization mediated by the combination of pY and GβGγ, we covalently linked pY peptides to supported membrane at a surface density of ∼5,000 pY/μm^2^ and then allowed farnesyl GβGγ to equilibrate into the membrane to density of ∼4,700 GβGγ/μm^2^. We quantified the membrane surface density of GβGγ at equilibrium using a combination of AF488-SNAP-GβGγ and dilute AF555-SNAP-GβGγ (0.0025%), to visualize the bulk and single molecule densities, respectively (**Figure 3A**). Comparing the bulk membrane absorption of Dy647-PI3Kβ in the presence of pY alone, we observed a 2-fold increase in membrane localization due to synergistic association with pY and GβGγ (**Figure 3B-3C**). Single molecule imaging experiments also showed a 1.9-fold increase in the membrane dwell time of Dy647-PI3Kβ in the presence of both pY and GβGγ (**Figure 3D**). Consistent with Dy647-PI3Kβ forming a complex with pY and GβGγ, we observed a 22% reduction in the average single particle displacement and a decrease in the diffusion coefficient due to ternary complex formation (**Figure 3E**).

**Figure 3.**
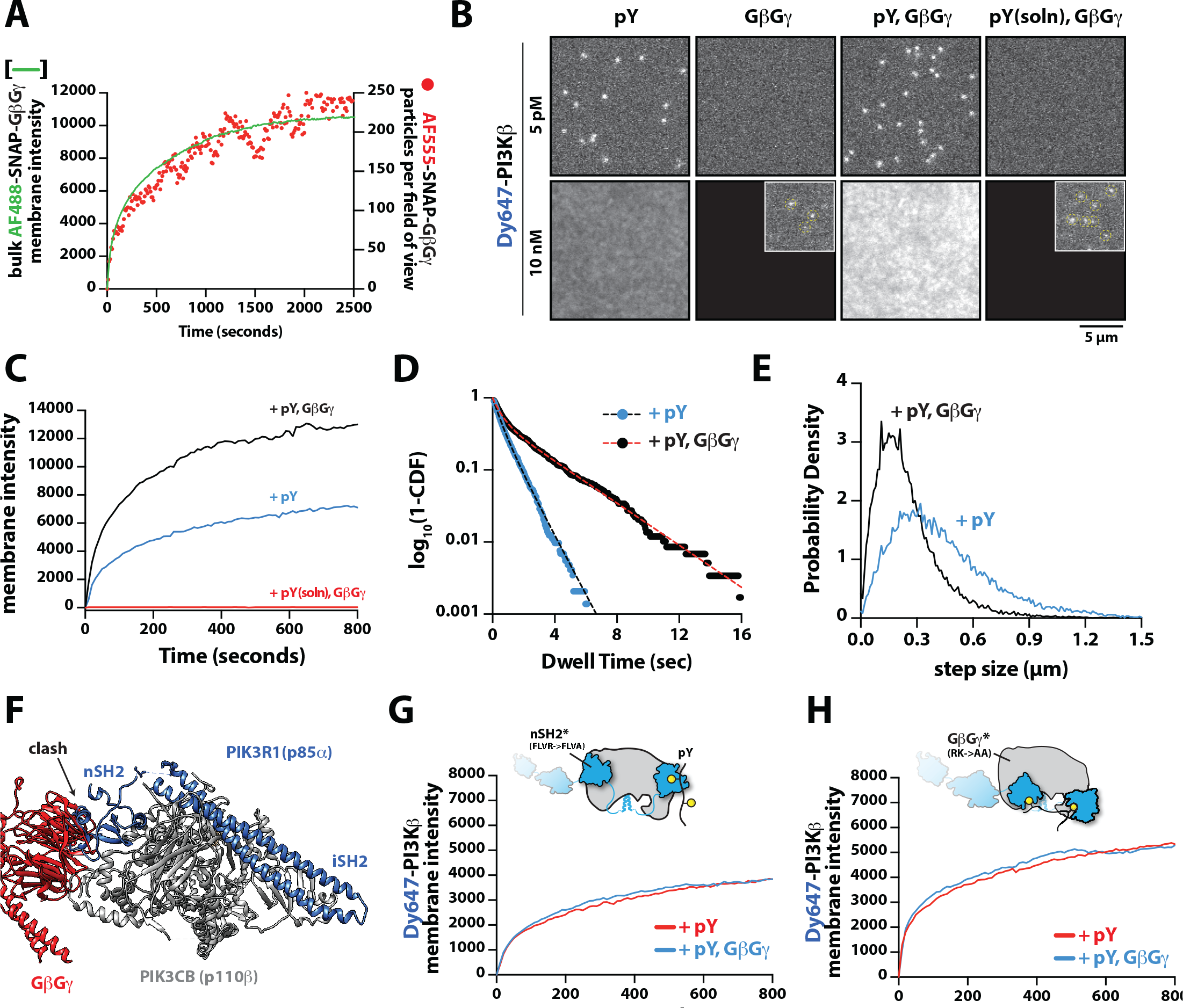
Mechanism controlling synergistic Dy647-PI3Kβ membrane binding by pY and GβGγ. **(A)** Kinetic trace showing the membrane absorption of 200 nM AF488-SNAP-GβGγ and AF555-SNAP-GβGγ (0.0025%) measured by TIRF-M. Single molecule densities of AF555-SNAP-GβGγ were calculated for each frame in a field of view of 3000 μm^2^. **(B)** Representative TIRF-M images showing the equilibrium membrane localization of 5 pM and 10 nM Dy647-PI3Kβ on membranes containing either pY, GβGγ, pY/GβGγ, or pY(solution)/GβGγ. The inset image (+GβGγ and +pY/GβGγ) shows low frequency single molecule binding events detected in the presence of 10 nM Dy647-PI3Kβ. Supported membranes were conjugated with 10 μM pY peptide (final surface density of ∼15,000 pY/μm^2^) and equilibrated with 200 nM farnesyl-GβGγ before adding Dy647-PI3Kβ. pY(solution) = 10 μM. **(C)** Bulk membrane recruitment dynamics of 10 nM Dy647-PI3Kβ measured in the presence of either pY alone, pY/GβGγ, or pY(solution)/GβGγ. pY(solution) = 10 μM. **(D)** Single molecule dwell time distributions measured in the presence of 5 pM Dy647-PI3Kβ on supported membranes containing pY alone (τ_1_=0.55±0.11s, τ_2_=1.44±0.56s, α=0.54, *N*=4698 particles, n=5 technical replicates) or pY/GβGγ (τ_1_=0.61±0.13s, τ_2_=3.09±0.27s, α=0.58, *N*=3421 particles, n=4 technical replicates). Alpha(α) represents the fraction of particles characterized by the time constant (τ_1_). **(E)** Step size distributions showing single molecule displacements measured in the presence of either pY alone (D1=0.34±0.04 μm^2^/sec, D2=1.02±0.07 μm^2^/sec, α=0.45) or pY/GβGγ (D1=0.23±0.03 μm^2^/sec, D2=0.88±0.08 μm^2^/sec, α=0.6); n=3-4 technical replicates from > 3000 tracked particles with 10,000-30,000 total displacements measured. Alpha(α) represents the fraction of particles characterized by the time constant (D1). **(F)** Combined model of the putative nSH2 and GβGγ binding sites on p110β. The p110β-GβGγ binding site is based on an Alphafold multimer model supported by previous HDX-MS and mutagenesis experiments. The orientation of the nSH2 is based on previous X-ray crystallographic data on PI3Kα (p110α-p85α, niSH2, PDB:3HHM) aligned to the structure of PI3Kβ (p110β-p85α, icSH2, PDB:2Y3A). **(G)** Bulk membrane recruitment dynamics of 10 nM Dy647-PI3Kβ, WT and nSH2(R358A), measured on membranes containing either pY or pY/GβGγ. **(H)** Bulk membrane recruitment dynamics of 10 nM Dy647-PI3Kβ, WT and GβGγ binding mutant, measured on membranes containing either pY or pY/GβGγ. **(A-H)** Membrane composition: 96% DOPC, 2% PI(4,5)P_2_, 2% MCC-PE.

Parallel to our experiments using membrane conjugated pY, we examined whether solution pY could promote Dy647-PI3Kβ localization to GβGγ-containing membranes. Based on the bulk membrane recruitment, solution pY did not strongly enhance membrane binding of Dy647-PI3Kβ on GβGγ-containing membranes on membranes that lack phosphatidylserine (PS) lipids (**Figure 3B**). Single molecule dwell analysis revealed few transient Dy647-PI3Kβ membrane interactions (inset **Figure 3A**) with a mean dwell time of 116 ms in the presence of GβGγ alone (**Figure 3 – figure supplement 1A**). The presence of 10 μM solution pY modestly increased the mean dwell time of Dy647-PI3Kβ to 136 ms on GβGγ containing membranes (**Figure 3 – figure supplement 1B**). When we measured the interaction between Dy647-PI3Kβ and GβGγ on membrane containing 20% PS, however, we observed a significant enhancement in Dy647-PI3Kβ localization when solution pY was added (**Figure 3 – figure supplement 1C-1D)**. This confirms that SH2 domain dependent inhibition of GβGγ binding can be relieved by solution pY, as previously reported (Hashem A. Dbouk et al. 2012). However, a high density of anionic lipids (i.e. PS) was required to facilitate the interaction between pY and GβGγ. This suggests that the affinity between PI3Kβ and GβGγ is relatively weak, which is consistent with previous structural biochemistry studies (Hashem A. Dbouk et al. 2012).

For PI3Kβ to engage GβGγ, it is hypothesized that the nSH2 domain must move out of the way from sterically occluding the GβGγ binding site. This model is supported by previous hydrogen deuterium exchange mass spectrometry (HDX-MS) experiments that could only detected interactions between GβGγ and PI3Kβ (p110β) when the nSH2 domain was either deleted or disengaged from the catalytic domain by a soluble RTK-derived pY peptide (Hashem A. Dbouk et al. 2012). We examined the putative interface of GβGγ bound to the p110β catalytic domain using AlphaFold multimer (Jumper et al. 2021; Evans et al. 2022; Varadi et al. 2022) which defined hα1 in the helical domain as the binding site. This result was consistent with previous mutagenesis and HDX-MS analysis of GβGγ binding to p110β (Hashem A. Dbouk et al. 2012). Comparing our model to previous X-ray crystallographic data of SH2 binding to either p110α and p110β (Zhang et al. 2011; Mandelker et al. 2009) suggested that the nSH2 domain sterically obstructs the GβGγ binding interface (**Figure 3F** and **Figure 3 – figure supplement 2**), with GβGγ activation only possible when the when the nSH2 dissociates from p110β interface. To test this hypothesis, we measured the membrane binding dynamics of Dy647-PI3Kβ nSH2(R358A) on membranes containing pY and GβGγ. Comparing the bulk membrane recruitment of these constructs revealed that the inability of the Dy647-PI3Kβ nSH2 domain to bind to pY peptides, made the kinase insensitive to synergistic membrane recruitment mediated by pY and GβGγ (**Figure 3G**). Similarly, the membrane association dynamics of Dy647-PI3Kβ nSH2(R358A), phenocopied a PI3Kβ (K532D, K533D) mutant that lacks the ability to engage GβGγ (**Figure 3H**).

### Rac1(GTP) and pY synergistically enhance PI3Kβ membrane localization

PI3Kβ is the only class IA PI3K that has been shown to interact with Rho-family GTPases, Rac1 and Cdc42 (Fritsch et al. 2013). Our membrane localization studies indicate however that Dy647-PI3Kβ does not strongly localize to membranes containing Rac1(GTP) alone (**Figure 1C-1D** and **Figure 1 – figure supplement 2B**). To determine whether membrane anchored pY peptides can facilitate interactions with Rac1(GTP), we visualized the localization of Dy647-PI3Kβ on membranes containing pY-Rac1(GDP) or pY-Rac1(GTP). Our experiments were designed to have the same pY surface density across conditions. By incorporating a small fraction of Cy3-Rac1 and Alexa488-pY into our Rac1-pY membrane coupling reaction we were able to visualize single membrane anchored proteins and calculate the membrane surface density of ∼4,000 Rac1/μm^2^ and ∼5,000 pY/ μm^2^ (**Figure 4A-4B**). Bulk localization to membranes containing either pY-Rac1(GDP) or pY-Rac1(GTP), revealed that active Rac1 could enhance Dy647-PI3Kβ localization 1.4-fold (**Figure 4C-4D**). Similarly, single molecule analysis revealed a 1.5-fold increase in the mean dwell time of Dy647-PI3Kβ in the presence of pY-Rac1(GTP) (**Figure 4E**). The average displacement of Dy647-PI3Kβ per frame (i.e. 52 ms) also decreased by 28% in the presence of pY and Rac1(GTP) (**Figure 4F**), consistent with the formation of a membrane bound PI3Kβ-pY-Rac1(GTP) ternary complex.

**Figure 4.**
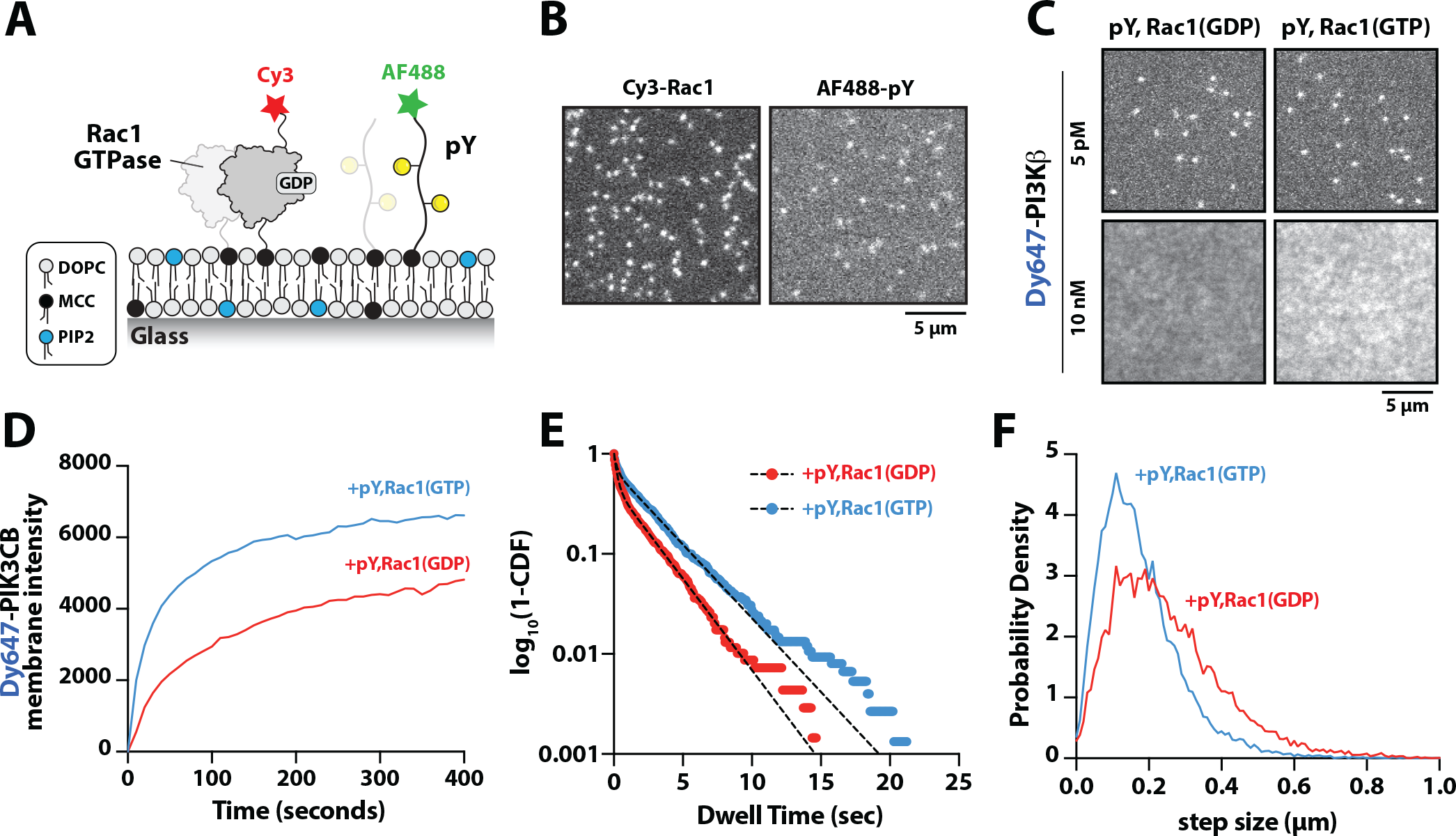
Membrane anchored pY peptides synergistically enhance Dy647-PI3Kβ membrane binding in the presence of Rac1(GTP) **(A)** Cartoon schematic showing membrane conjugation of Cy3-Rac1 and AF488-pY on membranes containing unlabeled Rac1 and pY. **(B)** Representative TIRF-M images showing localization of Cy3-Rac1 (1:10,000 dilution) and AF488-pY (1:30,000 dilution) after membrane conjugation in the presence of 30 μM Rac1 and 10 μM pY. Membrane surface density equals ∼4,000 Rac1/μm^2^ and ∼5,000 pY/μm^2^. **(C)** Representative TIRF-M images showing the equilibrium membrane localization of 5 pM and 10 nM Dy647-PI3Kβ measured in the presence of membranes containing either pY/Rac1(GDP) or pY/Rac1(GTP). **(D)** Bulk membrane recruitment dynamics of 10 nM Dy647-PI3Kβ measured in the presence of pY/Rac1(GDP) or pY/Rac1(GTP). **(E)** Single molecule dwell time distributions measured in the presence of 5 pM Dy647-PI3Kβ on supported membranes containing pY/Rac1(GDP) or pY/Rac1(GTP). **(F)** Step size distributions showing single molecule displacements from > 500 Dy647-PI3Kβ particles (>10,000 steps) in the presence of either pY/Rac1(GDP) or pY/Rac1(GTP). Membrane composition: 96% DOPC, 2% PI(4,5)P_2_, 2% MCC-PE.

### Rac1(GTP) and GβGγ stimulate PI3Kβ activity beyond enhancing membrane localization

Previous *in vitro* measurements of PI3Kβ activity have shown that solution pY stimulates lipid kinase activity (Zhang et al. 2011; Hashem A. Dbouk et al. 2012). Similar mechanisms of activation have been reported for other class IA kinases, including PI3Kα and PI3Kδ (Buckles et al. 2017; Burke et al. 2011; Dornan et al. 2017). Functioning in concert with pY peptides, GβGγ or Rho-family GTPase synergistically enhance PI3Kβ activity by a mechanism that remains unclear (Fritsch et al. 2013; Hashem A. Dbouk et al. 2012). Similarly, RTK derived peptides and H-Ras(GTP) have been shown to synergistically activate PI3Kα (Buckles et al. 2017; Siempelkamp et al. 2017; Yang et al. 2012). In the case of PI3Kβ, previous experiments have not determined whether synergistic activation by multiple signaling inputs results from an increase in membrane affinity (*K*_*D*_) or direct modulation of lipid kinase activity (*k*_*cat*_) through an allosteric mechanism. To determine how the lipid kinase activity of the PI3Kβ-pY complex is synergistically modulated by either GβGγ or Rho-family GTPases, we used TIRF-M to simultaneously visualize Dy647-PI3Kβ membrane localization and monitor PI(3,4,5)P_3_ production. To measure the kinetics of PI(3,4,5)P_3_ formation, we purified and fluorescently labeled the pleckstrin homology and Tec homology (PH-TH) domain derived from Bruton’s tyrosine kinase (Btk). We used a form of Btk containing a mutation that disrupts the peripheral PI(3,4,5)P_3_ lipid binding domain (Wang et al. 2015). This Btk mutant was previously shown to associate with a single PI(3,4,5)P_3_ head group and exhibits rapid membrane equilibration kinetics *in vitro* (Chung et al. 2019). Consistent with previous observations, Btk fused to SNAP-AF488 displayed high specificity and rapid membrane equilibration kinetics on SLBs containing PI(3,4,5)P_3_ (**Figure 5A-5B**). Compared to the PI(3,4,5) P_3_ lipid sensor derived from the Cytohesin/Grp1 PH domain (He et al. 2008; J. D. Knight et al. 2010), the Btk mutant sensor exhibited a faster association rate constant (*k*_*ON*_) and a more transient dwell time (1/ *k*_*OFF*_) making it ideal for kinetic analysis of PI3Kβ lipid kinase activity (**Figure 5 – figure supplement 1**). Using Btk-SNAP-AF488, we measured the production of PI(3,4,5)P_3_ lipids on SLBs by quantifying the time dependent recruitment in the presence of PI3Kβ. The change in Btk-SNAP-AF488 membrane fluorescence could be converted to the absolute number of PI(3,4,5) P_3_ lipids produced per μm^2^ to determine the catalytic efficiency per membrane bound Dy647-PI3Kβ.

**Figure 5.**
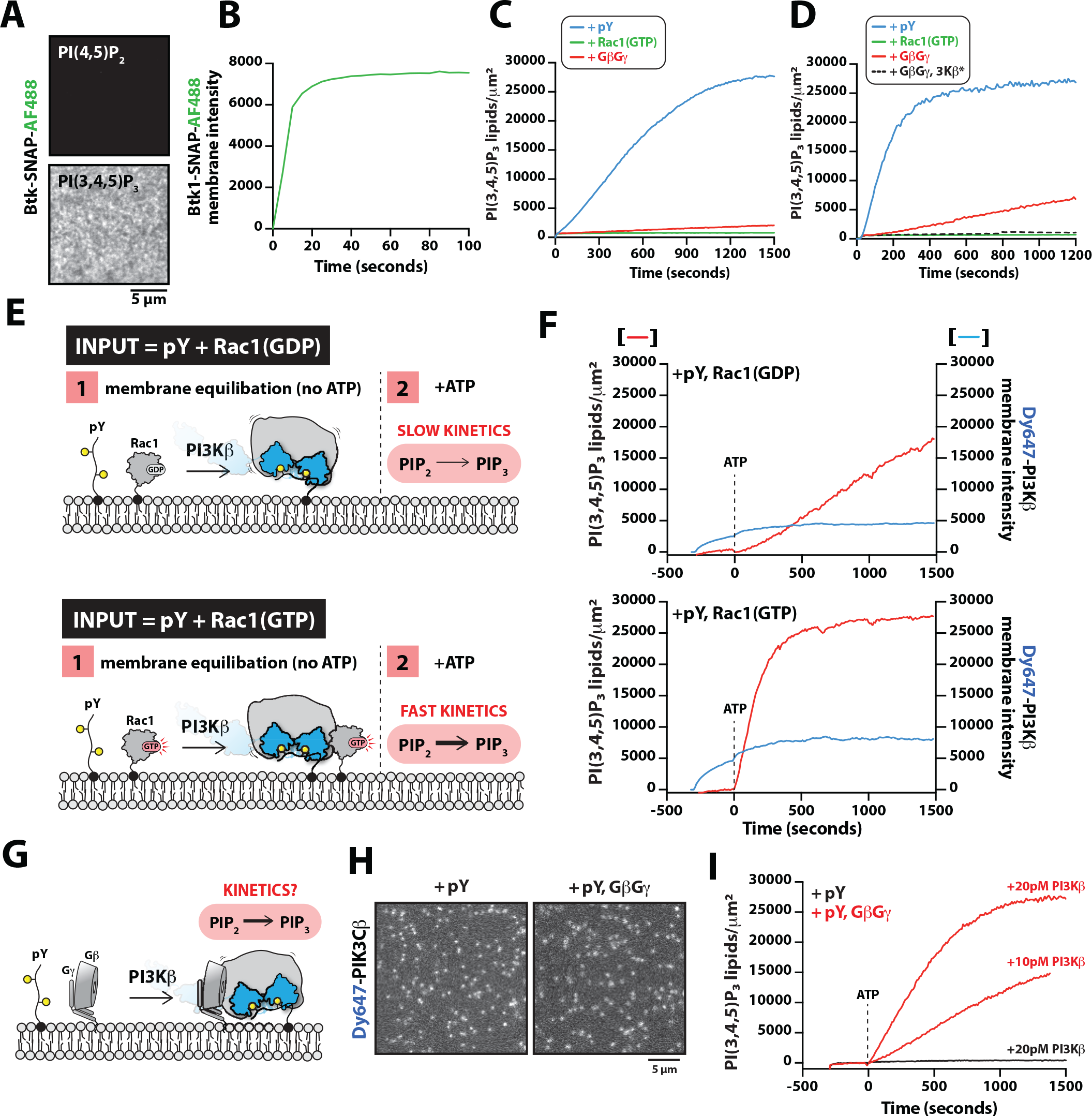
GβGγ and Rac1(GTP) stimulate PI3Kβ activity beyond enhancing localization on pY membranes. **(A)** Representative TIRF-M images showing localization of 20 nM Btk-SNAP-AF488 on SLBs containing either 2% PI(4,5)P_2_ or 2% PI(3,4,5)P_3_, plus 98% DOPC. **(B)** Bulk membrane recruitment kinetics of 20 nM Btk-SNAP-AF488 on an SLB measured by TIRF-M. **(C-D)** Kinetics of PI(3,4,5)P_3_ production measured in the presence of 10 nM Dy647-PI3Kβ and 1 mM ATP on SLBs with membrane anchored pY, Rac1(GTP), or GβGγ alone. Reactions in **(C)** were performed in the absence of PS lipids, while membranes in **(D)** contained 20% DOPS. **(E)** Cartoon schematic illustrating method for measuring Dy647-PI3Kβ activity in the presence of either pY/Rac1(GDP) or pY/ Rac1(GTP). Phase 1 of the reconstitution involves membrane equilibration of Dy647-PI3Kβ in the absence of ATP. During phase 2, 1 mM ATP is added to stimulate lipid kinase activity of Dy647-PI3Kβ. **(F)** Dual color TIRF-M imaging showing 2 nM Dy647-PI3Kβ localization and catalysis measured in the presence of 20nM Btk-SNAP-AF488. Dashed line represents the addition of 1 mM ATP to the reaction chamber. **(G)** Cartoon schematic showing experimental design for measuring synergistic binding and activation of Dy647-PI3Kβ in the presence of pY and GβGγ. **(H)** Representative single molecule TIRF-M images showing the localization of 20 pM Dy647-PI3Kβ in (G). **(I)** Kinetics of PI(3,4,5)P_3_ production monitored in the presence of 20 nM Btk-SNAP-AF488 and 20 pM Dy647-PI3Kβ. Membrane contained either pY or pY/GβGγ. **(B, C, F, H, I)** Membrane composition: 96% DOPC, 2% PI(4,5)P_2_, 2% MCC-PE. **(D)** Membrane composition: 76% DOPC, 20% DOPS, 2% PI(4,5)P_2_, 2% MCC-PE. All kinetic measurements of PI(3,4,5)P_3_ production were performed in the presence of 20 nM Btk-SNAP-AF488.

While our findings provide a mechanism for enhanced PI3Kβ membrane localization in the presence of either pY-Rac1(GTP) or pY-GβGγ, these results did not reveal the mechanism controlling synergistic activation of lipid kinase activity. To probe if synergistic activation results from enhanced membrane localization or allosteric modulation of PI3Kβ, we first examined how well the pY peptide stimulates PI3Kβ lipid kinase activity on SLBs. In the absence of pY peptides, PI3Kβ did not catalyze the production of PI(3,4,5)P_3_ lipids, while the addition of 10 μM pY in solution resulted in a subtle but detectable increase in PI3Kβ lipid kinase activity (**Figure 5 – figure supplement 2A-2C**). By contrast, covalent conjugation of pY peptides to supported lipid bilayers increased the rate of PI(3,4,5)P_3_ production by 207-fold (**Figure 5 – figure supplement 2A-2C**). The observed difference in kinetics was consistent with robust membrane recruitment of Dy647-PI3Kβ requiring membrane tethered pY peptides. Comparing the lipid kinase activity of PI3Kβ measured in the presence of individual signaling inputs – pY, Rac1(GTP), and GβGγ – revealed that pY alone provided the strongest stimulation (**Figure 5C**). This was largely due to the dramatic enhancement in membrane recruitment caused by membrane tethered pY peptides. Incorporation of 20% DOPS lipids in the SLBs raised of the overall activity of PI3Kβ across all conditions but the general trend for signaling input preference was preserved (**Figure 5D**). Similar to previous observations (Hashem A. Dbouk et al. 2012), we found that GβGγ alone could stimulate PI3Kβ lipid kinase activity, without strongly enhancing membrane PI3Kβ localization (**Figure 1 – figure supplement 2**). By contrast, the PI3Kβ (K532D, K533D) mutant that is unable to interact with GβGγ was insensitive to GβGγ-mediated activation (**Figure 5D**).

Next, we sought to assess if the combination of pY and Rac1(GTP) could synergistically stimulate PI3Kβ activity beyond the expected increase due to the enhanced membrane localization of the PI3Kβ-pY-Rac1(GTP) complex. To decipher the mechanism of synergistic activation, we performed two-phase experiments that accounted for both the total amount of membrane localized Dy647-PI3Kβ and the corresponding kinetics of PI(3,4,5)P_3_ generation. In phase 1 of our experiments, Dy647-PI3Kβ was flowed over SLBs and allowed to equilibrate with either pY-Rac1(GDP) or pY-Rac1(GTP) in the absence of ATP (**Figure 5E-5F**). This resulted in a 1.8-fold increase in Dy647-PI3Kβ localization mediated by the combination of membrane tethered Rac1(GTP) and pY, compared to pY membranes alone. Following membrane equilibration of Dy647-PI3Kβ, phase 2 was initiated by adding 1 mM ATP to the reaction chamber to stimulate lipid kinase activity. We found that the addition of ATP did not alter the bulk localization of Dy647-PI3Kβ, though the kinase was in dynamic equilibrium between the solution and membrane. Conducting experiments in this manner allowed us to measure activation by inputs while removing uncertainty from differential Dy647-PI3Kβ association with various signaling inputs. After accounting for the 1.8-fold difference in Dy647-PI3Kβ membrane localization comparing pY-Rac1(GDP) and pY-Rac1(GTP) membranes, we calculated a 4.3-fold increase in PI3Kβ activity that was dependent on Rac1(GTP).

Using the two-phase kinase assay described above, we next examined how pY and GβGγ synergistically activate PI3Kβ (**Figure 5G**). In our pilot experiments, we immediately observed more robust activation of PI3Kβ in the presence of pY-GβGγ, compared to pY-Rac1(GTP). To accurately measure the rapid kinetics of PI(3,4,5)P_3_ generation on SLBs we had to use a 100-fold lower concentration of Dy647-PI3Kβ. Under these conditions, single membrane bound Dy647-PI3Kβ molecules could be spatially resolved, which allowed us to measure the catalytic efficiency per PI3Kβ (**Figure 5H**). Comparing the activity of Dy647-PI3Kβ on membranes with pY alone or pY-GβGγ, we observed a 22-fold increase in catalytic efficiency comparing the PI3Kβ-pY and PI3Kβ-pY-GβGγ complexes (**Figure 5I**). Synergistic activation was dependent on the direct interaction between PI3Kβ and GβGγ (**Figure 5 – figure supplement 3A**). By varying the density of membrane anchored GβGγ we determined that maximum synergistic activation occurred when GβGγ was present at a density greater than ∼2,400 molecules/μm^2^ (**Figure 5 – figure supplement 2D**). Similar levels of PI3Kβ activity were measured in the ATP spike experiments when 20% DOPS was incorporated in the supported membranes (**Figure 5 – figure supplement 2E**). Based on membrane bound density of ∼0.2 Dy647-PI3Kβ molecules per μm^2^, we calculate a *k*_*cat*_ of 57 PI(3,4,5)P_3_ lipids/sec•PI3Kβ on pY-GβGγ containing membranes. By contrast, the Dy647-PI3Kβ-pY complex had a *k*_*cat*_ of ∼3 PI(3,4,5)P_3_ lipids/sec•PI3Kβ.

## DISCUSSION

### Prioritization of signaling inputs

The exact mechanisms that regulate how PI3Kβ prioritizes interactions with signaling input, such as pY, Rac1(GTP), and GβGγ remains unclear. To fill this gap in knowledge, we directly visualized the membrane association and dissociation dynamics of fluorescently labeled PI3Kβ on supported lipid bilayers using single molecule TIRF microscopy. This is the first study to reconstitute membrane localization and activation of a class 1A PI3K using multiple signaling inputs that are all membrane tethered in a physiologically relevant configuration. Previous experiments have relied exclusively on phosphotyrosine peptides (pY) presented in solution to activate PI3Kα, PI3Kβ, or PI3Kδ (Zhang et al. 2011; Dornan et al. 2017; Hashem A. Dbouk et al. 2012). However, pY peptides are derived from the cytoplasmic domains of transmembrane receptors, such as receptor tyrosine kinases (RTKs), which reside in the plasma membrane (Lemmon and Schlessinger 2010). Although pY peptides in solution can disrupt the inhibitory contacts between the regulatory and catalytic subunits of class 1A PI3Ks (Zhang et al. 2011; Yu et al. 1998), they do not robustly localize PI3Ks to membranes when they exist in solution. When conjugated to a SLB we find that pY peptides strongly localize PI3Kβ and relieve auto-inhibition, while membranes containing either Rac1(GTP) or GβGγ alone are unable to robustly localize PI3Kβ. We observed this prioritization of signaling input interactions over a range of membrane compositions that contained physiologically relevant densities of anionic lipids, such as 20% phosphatidylserine and 2% PI(4,5)P_2_. Although a small fraction of PI3Kβ may transiently adopt a conformation that is compatible with direct Rac1(GTP) or GβGγ association in the absence of pY, these events are rare and do not represent the most probable pathway for controlling initial PI3Kβ membrane docking. Given the complex of the cellular plasma membrane lipid composition, our experimental system uses a relatively simple mixture of lipids that maximize membrane fluidity and minimize surface defects that can promote non-specific molecular interactions. By this approach, our study provides clarity concerning the strength of various protein-protein interactions that regulate PI3Kβ membrane localization and activity.

Based on our single molecule dwell time and diffusion analysis, Dy647-PI3Kβ can cooperatively bind to one doubly phosphorylated peptide derived from the PDGF receptor. Supporting this model, Dy647-PI3Kβ with a mutated nSH2 or cSH2 domain that eliminates pY binding, still displayed membrane diffusivity indistinguishable from wild-type PI3Kβ. The diffusion coefficient of membrane bound pY-PI3Kβ complexes also did not significantly change over a broad range of pY membrane surface densities that span 3 orders of magnitude. Given that diffusivity of peripheral membrane binding proteins is strongly correlated with the valency of membrane interactions (Ziemba and Falke 2013; Hansen et al. 2022), we expected to observe a decrease in Dy647-PI3Kβ diffusion with increasing membrane surface densities of pY. Instead, our data suggests that the vast majority of PI3Kβ molecules engage a single pY peptide, rather than binding one tyrosine phosphorylated residue on two separate pY peptides. While no structural studies have shown how exactly the tandem SH2 domains of PI3K (p85α) simultaneously bind to a doubly phosphorylated pY peptide, the interactions likely resemble the mechanism reported for ZAP-70 (ζ-chain of T-cell Receptor Associated Protein Kinase 70). The tandem SH2 domains of ZAP-70 can bind to a doubly phosphorylated ζ-chain derived from the TCR with only 11 amino acids spacing between the two tyrosine phosphorylation sites (Hatada et al. 1995). In the case of our PDGFR derived pY peptide that binds p85α, 10 amino acids separate the two tyrosine phosphorylated residues.

Following the engagement of a pY peptide, we find that PI3Kβ can then associate with membrane anchored Rac1(GTP) or GβGγ. We detected the formation of PI3Kβ-pY-Rac1(GTP) and PI3Kβ-pY-GβGγ complexes based on the following criteria: (1) increase in Dy647-PI3Kβ bulk membrane recruitment, (2) increase in single molecule dwell time, and (3) a decrease in membrane diffusivity. Consistent with Dy647-PI3Kβ having a weak affinity for GβGγ, pY peptides in solution were unable to strongly localize Dy647-PI3Kβ to SLBs containing membrane anchored GβGγ. This is in agreement with HDX-MS data showing that the p110β-GβGγ interaction can only be detected using a GβGγ-p85α(icSH2) chimeric fusion or pre-activating PI3Kβ with solution pY (Hashem A. Dbouk et al. 2012). Using AlphaFold Multimer (Evans et al. 2022; Jumper et al. 2021), we created a model that illustrates how the p85α(nSH2) domain is predicted to sterically block GβGγ binding to p110β. This model was validated by comparing the AlphaFold Multimer model to previous reported HDX-MS (H. A. Dbouk et al. 2010) and X-ray crystallography data (Zhang et al. 2011). Further supporting this model, we found that the Dy647-PI3Kβ nSH2(R358A) mutant tethered to membrane conjugated pY peptide was unable to engage membrane anchored GβGγ. Membrane targeting of PI3Kβ by pY was required to relieve nSH2 mediated autoinhibition and expose the GβGγ binding site. Recruitment by membrane tethered pY also reduces the translational and rotational entropy of PI3Kβ, which facilitates PI3Kβ-pY-GβGγ complex formation. We observed a similar mechanism of synergistic PI3Kβ localization on SLBs containing pY and Rac1(GTP). This is consistent with single molecule studies investigating synergistic activation of PI3Kα in the presence of H-Ras(GTP) and solution pY peptide (Buckles et al. 2017). It remains unclear how p85α mediated inhibition controls the association dynamics between PI3Kβ and Rac1(GTP).

### Mechanism of synergistic activation

Previous characterization of PI3Kβ lipid kinase activity has utilized solution-based assays to measure P(3,4,5)P_3_ production. These solution-based measurements lack spatial information concerning the mechanism of PI3Kβ membrane recruitment and activation. Our ability to simultaneously visualize PI3Kβ membrane localization and P(3,4,5)P_3_ production is critical for determining which regulatory factors directly modulate the catalytic efficiency of PI3Kβ. In the case of PI3Kα, the enhanced membrane recruitment model has been used to explain the synergistic activation mediated by pY and Ras(GTP) (Buckles et al. 2017). In other words, the PI3Kα-pY-Ras complex is more robustly localized to membranes compared to the PI3Kα-pY and PI3Kα-Ras complexes, which results in a larger total catalytic output for the system. Although the Ras binding domain (RBD) of PI3Kα and PI3Kβ are conserved, these kinases interact with distinct Ras superfamily GTPases (Fritsch et al. 2013). Therefore, it’s possible that PI3Kα and PI3Kβ display different mechanisms of synergistic activation, which could explain their non-overlapping roles in cell signaling.

Studies of PI3Kβ mouse knock-in mutations in primary macrophages and neutrophils have shown that robust PI3Kβ activation requires coincident activation through the RTK and GPCR signaling pathways (Houslay et al. 2016). This response most strongly depends on the ability of PI3Kβ to bind GβGγ and to a lesser extent Rac1/Cdc42 (Houslay et al. 2016). A similar mechanism of synergistic activation has been reported for PI3Kγ in the presence of H-Ras(GTP) and GβGγ (Suire et al. 2012). Although mutational studies have nicely demonstrated the pathways that drive synergistic of PI3Kγ and PI3Kβ activation in cells, signaling network crosstalk and redundancy limits our mechanistic understanding of how these kinases prioritizes signaling inputs and the exact mechanism for driving PI(3,4,5)P_3_ production. Based on our single molecule membrane binding experiments, auto-inhibited PI3Kβ does not strongly associate with either Rac1(GTP) or GβGγ in the absence of pY peptides. We found that PI3Kβ kinase activity is also relatively insensitive to either Rac1(GTP) or GβGγ alone but can stimulate some PI3Kβ activity when 20% PS lipids is incorporated in our supported lipid bilayers. Importantly, the prioritization of signal input localization strengths was similar across all membrane compositions tested. Previous studies that showed Rho-GTPases (Fritsch et al. 2013) and GβGγ (Katada et al. 1999; Hashem A. Dbouk et al. 2012; Maier, Babich, and Nürnberg 1999) can individually activate PI3Kβ used more complex lipid mixtures that incorporate sphingomyelin, cholesterol, and phosphatidylethanolamine to mimic the cellular plasma membrane composition. In the future, a more comprehensive analysis will be required to map the relationship between PI3Kβ activity, membrane localization, and lipid composition.

In our single molecule TIRF experiments, we find that the pY peptide is the only factor that robustly localizes PI3Kβ to supported membranes in an autonomous manner. However, the pY-PI3Kβ complex displays weak lipid kinase activity (*k*_*cat*_ = ∼3 PI(3,4,5)P_3_ lipids/sec•PI3Kβ). This is consistent with cellular measurements showing that RTK activation by insulin (Z. A. Knight et al. 2006), PDGF (Guillermet-Guibert et al. 2008), or EGF (Ciraolo et al. 2008) show little PI3Kβ dependence for PI(3,4,5)P_3_ production. Although the dominant role of PI3Kα in controlling PI(3,4,5)P_3_ production downstream of RTKs can mask the contribution from PI3Kβ in some cell types, these results highlight the need for PI3Kβ to be synergistically activated. When we measured the kinetics of lipid phosphorylation for PI3Kβ-pY-Rac1(GTP) and PI3Kβ-pY-GβGγ complexes we observed synergistic activation beyond simply enhancing PI3Kβ membrane localization. After accounting for the ∼1.8-fold increase in membrane localization between PI3Kβ-pY-Rac1(GTP) and PI3Kβ-pY-Rac1(GDP), we calculated a 4.3-fold increase in *k*_*cat*_ (13 PI(3,4,5)P_3_ lipids/sec•PI3Kβ) that was dependent on engaging Rac1(GTP). Comparing the kinase activity of PI3Kβ-pY and PI3Kβ-pY-GβGγ complexes that are present at the same membrane surface density (∼0.2 PI3Kβ/ μm^2^) revealed a 22-fold increase in *k*_*cat*_ mediated by the GβGγ interaction. Together, these results indicate that PI3Kβ-pY complex association with either Rac1(GTP) or GβGγ allosterically modulates PI3Kβ, making it more catalytically efficient.

### Mechanisms controlling cellular activation of PI3Kβ

Studies of PI3K activation by pY peptides have mostly been performed using peptides derived from the IRS-1 (Insulin Receptor Substrate 1) and the EGFR/ PDGF receptors (Backer et al. 1992; Fantl, Martin, and Turck 1992). As a result, we still have not defined the broad specificity p85α has for tyrosine phosphorylated peptides. Biochemistry studies indicate that the nSH2 and cSH2 domains of p85α robustly bind pY residues with a methionine in the +3 position (pYXXM) (Breeze et al. 1996; Nolte et al. 1996; Backer et al. 1992; Fantl, Martin, and Turck 1992). The p85α subunit is also predicted to interact with the broad repertoire of receptors that contain immunoreceptor tyrosine-based activation motifs (ITAMs) baring the pYXX(L/I) motif (Reth 1989; Osman et al. 1996; Zenner et al. 1996; Love and Hayes 2010). Based on RNA seq data, human neutrophils express at least six different Fc receptors (FcRs) that all contain phosphorylated ITAMs that can potentially facilitate membrane localization of class 1A PI3Ks (Rincón, Rocha-Gregg, and Collins 2018).

A variety of human diseases result from the overexpression of RTKs, especially the epithelial growth factor receptor (EGFR) (Sauter et al. 1996). When the cellular plasma membrane contains densities of EGFR greater than 2000 receptors/μm^2^, trans-autophosphorylation and activation can occur in a EGF-independent manner (Endres et al. 2013). Receptor membrane surface densities above the threshold needed for spontaneous receptor trans-autophosphorylation have been observed in many cancer cells (Haigler et al. 1978). In these disease states, PI3K is expected to localize to the plasma membrane in the absence of ligand induced RTK or GPCR signaling. The slow rate of PI(3,4,5)P_3_ production we measured for the membrane tethered pY-PI3Kβ complex suggests that PI(3,4,5)P_3_ levels are not likely to rise above the global inhibition imposed by lipid phosphatases until synergistic activation of PI3Kβ by RTKs and GPCRs. However, loss of PTEN in some cancers (Jia et al. 2008) could produce an elevated level of PI(3,4,5)P_3_ due to PI3Kβ being constitutively membrane localized via ligand independent trans-autophosphorylation of RTKs.

## ACKNOWLEDGMENTS

We thank John Burke (University of Victoria) for assistance generating the AlphaFold2 Multimer model of PI3Kβ bound to GβGγ. We thank Grace Waddell for preliminary characterization of PI3Kβ. We thank Colin Hawkinson for assistance with protein purification. We thank Jean Chung (Colorado State, Fort Collins) and Orion Weiner (University of California at San Francisco) for plasmids encoding Btk and P-Rex1 plasmids, respectively.

## AUTHOR CONTRIBUTIONS

Resources: B.R.D, G.M.B, S.E.P, S.D.H.

Experiments and investigation: B.R.D, N.E.W., G.M.B, S.E.P, S.D.H.

Data Analysis: B.R.D, N.E.W., S.E.P., S.D.H.

Conceptualization: B.R.D, N.E.W., S.D.H. Interpretation: B.R.D, N.E.W., S.D.H.

Data curation: B.R.D, N.E.W., S.D.H.

Writing – Review and editing: B.R.D, N.E.W., G.M.B, S.E.P, S.D.H.

Writing – Original draft: S.D.H. Supervision: S.D.H.

Project administration: S.D.H. Funding acquisition: S.D.H.

## DATA AVAILABILITY

All the information needed for interpretation of the data is presented in the manuscript or the supplemental material. Plasmids related to this work are available upon request.

## MATERIALS & METHODS

### Molecular Biology

The following genes were used as templates for PCR to clone plasmids used for recombinant protein expression: *PIK3CB* (human 1-1070aa; Uniprot Accession #P42338), *PIK3R1* (human 1-724aa; Uniprot Accession #P27986), *PIK3CG* (mouse 1-1102aa; Uniprot Accession #Q9JHG7), *PIK3R5* (mouse 1-871aa; Uniprot Accession #Q5SW28), *RAC1* (human 1-192aa; Uniprot Accession #P63000), *CYTH3*/Grp1 (human 1-400aa; Uniprot Accession #O43739), *BTK* (bovine 1-659aa; Uniprot Accession #Q3ZC95), neutrophil cytosol factor 2 (*NCF2*, human 1-526aa; Uniprot Accession #P19878, referred to as p67/phox), *PREX1* (human 1-1659aa; Uniprot Accession #Q8TCU6), *GNB1/GBB1* (Gβ_1_, bovine 1-340aa; Uniprot Accession #P62871), *GNG2/GBG2* (Gγ_2_, bovine 1-71aa; Uniprot Accession #P63212). The following plasmids were purchased as cDNA clones from Horizon (PerkinElmer), formerly known as Open Biosystems and Dharmacon: mouse *PIK3CG* (clone #BC051246, cat #MMM1013-202770664) and mouse *PIK3R5* (clone #BC128076, cat #MMM1013-211693360), human *PIK3R1* (clone #30528412, cat #MHS6278-202806334), human *CYTH3*/Grp1 (clone #4811560, cat #MHS6278-202806616). Genes encoding bovine Gβ_1_ and Gγ_2_ were derived from the following plasmids: YFP-Gβ_1_ (Addgene plasmid # 36397) and YFP-Gγ_2_ (Addgene plasmid # 36102). These Gβ_1_ and Gγ_2_ containing plasmids were kindly provided to Addgene by Narasimhan Gautam (Saini et al. 2007). In this study, we used a previously described mutant form of Btk with mutations in the peripheral PI(3,4,5)P_3_ binding site (R49S/K52S) (Chung et al. 2019; Wang et al. 2015). The Btk peripheral site mutant was PCR amplified using a plasmid provided by Jean Chung (Colorado State, Fort Collins) that contained the following coding sequence: his6-SUMO-Btk(PH-TH, R49S/K52S)-EGFP. The nSH2 biosensor was derived from human *PIK3R1*. The gene encoding human *PREX1* was provided by Orion Weiner (University of California San Francisco). Refer to supplemental text to see exact peptide sequence of every protein purified in this study. The following mutations were introduced into either the *PIK3CB* (p110β) or *PIK3R1* (p85α) genes using site-directed mutagenesis: p85α nSH2 (R358A, FLVR->FLVA), p85α cSH2 (R649A, FLVR->FLVA), p110β GβGγ mutant (K532D/K533D). For cloning, genes were PCR amplified using AccuPrime *Pfx* master mix (ThermoFisher, Cat#12344040) and combined with a restriction digested plasmids using Gibson assembly (Gibson et al. 2009). Refer to supplemental text for a complete list of plasmids used in this study. Information about the specific peptide sequences for recombinantly expressed and purified proteins is organized in the supplemental information document. The complete open reading frame of all vectors used in this study were sequenced to ensure the plasmids lacked deleterious mutations.

### BACMID and baculovirus production

We generated BACMID DNA as previously described (Hansen et al. 2019). FASTBac1 plasmids containing our gene of interested were transformed into DH10 Bac cells and plated on LB agar media containing 50 μg/ mL kanamycin, 10 μg/mL tetracycline, 7 μg/mL gentamycin, 40 μg/mL X-GAL, and 40 μg/mL IPTG. Plated cells were incubated for 2-3 days at 37ºC before positive clones were isolated based on blue-white colony selection. White colonies were inoculated into 5mL of TPM containing 50 μg/mL kanamycin, 10 μg/mL tetracycline, 7 μg/ mL gentamycin and grown overnight at 37ºC. To purify the BACMID DNA, we first pelleted the cultures via centrifugation, then re-suspended the pellet in 300 μL of buffer containing 50 mM Tris [pH 8.0], 10 mM EDTA, 100 μg/mL RNase A. We lysed bacteria via addition of 300 μL of buffer containing 200 mM NaOH, 1% SDS before neutralization with 300 μL of 4.2 M Guanidine HCl, 0.9 M KOAc [pH 4.8]. We then centrifuged the sample at 23ºC for 10 minutes at 14,000 x g. Supernatant containing the BACMID DNA was combined with 700 μL 100% isopropanol and spun for 10-minute at 14,000 x g. The DNA pellets were washed twice with 70% ethanol (200 μL and 50 μL) and centrifuged. The ethanol was removed by vacuum aspiration and the final DNA pellet was dried in a biosafety hood. Finally, we solubilized the BACMID DNA in 50-100 μL of sterile filtered MilliQ water. A Nanodrop was used to quantify the total DNA concentration. BACMID DNA was be stored at -20ºC or used immediately for higher transfection efficiency. Baculovirus was generated as previously described. In brief, we incubated 5-7 μg of BACMID DNA with 4 μL Fugene (Thermo Fisher, Cat# 10362100) in 250 μL of Opti-MEM serum free media for 30 minutes at 23ºC. The DNA-Fugene mixture was then added to a Corning 6-well plastic dish (Cat# 07-200-80) containing 1 x 10^6^ *Spodoptera frugiperda* (Sf9) insect cells in 2 mL of ESF 921 Serum-Free Insect Cell Culture media (Expression Systems, Cat# 96-001, Davis, CA.). 4-5 days following the initial transfection, we harvested and centrifuged the viral supernatant (called “P0”). P0 was used to generate a P1 titer by infecting 7 x 10^6^ *Sf9* cells plated in a 10 cm tissue culture grade petri dish containing 10 mL of ESF 921 media and 2% Fetal Bovine serum (Seradigm, Cat# 1500-500, Lot# 176B14). We harvested and centrifuged the P1 titer after 4 days of transfection. The P1 titer was expanded at a concentration of 1% vol/vol of P1 titer into a 100 mL Sf9 cell culture grown to a density of 1.25-1.5 x 10^6^ cells/mL in a sterile 250 mL polycarbonate Erlenmeyer flask with vented cap (Corning, #431144). The P2 titer (viral supernatant) was harvested, centrifuged, and 0.22 μm filtered in 150 mL filter-top bottle (Corning, polyethersulfone (PES), Cat#431153). We used this P2 titer for protein expression in High 5 cells grown in ESF 921 Serum-Free Insect Cell Culture media (0% FBS) at a final baculovirus concentration of ∼2% vol/vol. All our media contained 1x concentration of Antibiotic-Antimycotic (Gibco/Invitrogen, Cat#15240-062).

### Protein purification

#### PI3Kβ and PI3Kγ

Genes encoding human his6-TEV-PIK3CB (1-1070aa) and ybbr-PIK3R1 (1-724aa) were cloned into a modified FastBac1 dual expression vector containing tandem polyhedrin (pH) promoters. Genes encoding mouse his6-TEV-PIK3CG (1-1102aa) and mouse his6-TEV-ybbr-PIK3R5 (1-871aa) were expressed from separate FastBac1 vectors under the polyhedrin (pH) promoters. For protein expression, high titer baculovirus was generated by transfecting 1 x 10^6^ *Spodoptera frugiperda* (Sf9) with 0.75-1μg of BACMID DNA as previously described (Hansen et al. 2019). After two rounds of baculovirus amplification and protein test expression, 2 x 10^6^ cells/mL High 5 cells were infected with 2% vol/vol PI3Kβ (PIK3CB/PIK3R1) or 2% vol/vol PI3Kγ (PIK3CG/PIK3R5) baculovirus and grown at 27ºC in ESF 921 Serum-Free Insect Cell Culture media (Expression Systems, Cat# 96-001) for 48 hours. High 5 cells were harvested by centrifugation and washed with 1x PBS [pH 7.2] and centrifuged again. Final cell pellets were resuspending in an equal volume of 1x PBS [pH 7.2] buffer containing 10% glycerol and 2x protease inhibitor cocktail (Sigma, Cat# P5726) before being stored in the -80ºC freezer. For protein purification, frozen cell pellets from 4 liters of cell culture were lysed by Dounce homogenization into buffer containing 50 mM Na_2_HPO_4_ [pH 8.0], 10 mM imidazole, 400 mM NaCl, 5% glycerol, 2 mM PMSF, 5 mM BME, 100 μg/mL DNase, 1x protease inhibitor cocktail (Sigma, Cat# P5726). Lysate was centrifuged at 35,000 rpm (140,000 x *g*) for 60 minutes under vacuum in a Beckman centrifuge using a Ti-45 rotor at 4ºC. Lysate was batch bound to 5 mL of Ni-NTA Agarose (Qiagen, Cat# 30230) resin for 90 minutes stirring in a beaker at 4ºC. Resin was washed with buffer containing 50 mM Na_2_HPO_4_ [pH 8.0], 30 mM imidazole, 400 mM NaCl, and 5 mM BME. Protein was eluted from NiNTA resin with wash buffer containing 500 mM imidazole. The his6-TEV-PIK3CB/ybbr-PIK3R1 complex was then desalted on a G25 Sephadex column in buffer containing 20 mM Tris [pH 8.0], 100 mM NaCl, 1 mM DTT. Peak fractions were pooled and loaded onto a Heparin anion exchange column equilibrated in 20 mM Tris [pH 8.0], 100 mM NaCl, 1 mM DTT buffer. Proteins were resolved over a 10-100% linear gradient (0.1-1 M NaCl) at 2 mL/min flow rate over 20 minutes. Peak fractions were pooled and supplemented with 10% glycerol, 0.05% CHAPS, and 200 μg/mL his6-TEV(S291V) protease. The his6-TEV-PIK3CB/ybbr-PIK3R1 complex was incubated overnight at 4ºC with TEV protease to remove the affinity tag. The TEV protease cleaved PIK3CB/ybbr-PIK3R1 complex was separated on a Superdex 200 size exclusion column (GE Healthcare, Cat# 17-5174-01) equilibrated with 20 mM Tris [pH 8.0], 150 mM NaCl, 10% glycerol, 1 mM TCEP, 0.05% CHAPS. Peak fractions were concentrated in a 50 kDa MWCO Amicon centrifuge tube and snap frozen at a final concentration of 10 μM using liquid nitrogen. This protein is referred to as PI3Kβ throughout the manuscript. The same protocol was followed to purify mouse PI3Kγ (PIK3CG/ybbr-PIK3R5) and the various PI3Kβ mutants reported in this study.

#### Rac1

The gene encoding human Rac1 were expressed in BL21 (DE3) bacteria as his10-SUMO3-(Gly)_5_ fusion proteins. Bacteria were grown at 37°C in 4L of Terrific Broth for two hours or until OD_600_ = 0.8. Cultures were shifted to 18°C for 1 hour then induced with 0.1 mM IPTG. Expression was allowed to continue for 20 hours before harvesting. Cells were lysed into 50 mM Na_2_HPO_4_ [pH 8.0], 400 mM NaCl, 0.4 mM BME, 1 mM PMSF, 100 μg/mL DNase using a microfluidizer. Lysate was centrifuged at 16,000 rpm (35000 x g) for 60 minutes in a Beckman JA-20 rotor at 4ºC. Lysate was circulated over 5 mL HiTrap Chelating column (GE Healthcare, Cat# 17-0409-01) loaded with CoCl_2_. Bound protein was eluted at a flow rate of 4mL/min into 50 mM Na_2_HPO_4_ [pH 8.0], 400 mM NaCl, 500mM imidazole. Peak fractions were pooled and combined with SUMO protease (SenP2) at a final concentration of 50μg/mL and dialyzed against 4 liters of buffer containing 20 mM Tris [pH 8.0], 250 mM NaCl, 10% Glycerol, 1 mM MgCl_2_, 0.4mM BME. Dialysate containing SUMO cleaved protein was recirculated for 2 hours over a 5 mL HiTrap Chelating column. Flowthrough containing (Gly)_5_-Rac1 was concentrated in a 5 MWCO Vivaspin 20 before being loaded on a 124 mL Superdex 75 column equilibrated in 20 mM Tris [pH 8.0], 150 mM NaCl, 10% Glycerol, 1 mM TCEP, 1 mM MgCl_2_. Peak fractions containing (Gly)_5_-Rac1 were pooled and concentrated to a concentration of 400-500 μM (∼10 mg/mL) and snap frozen with liquid nitrogen and stored at -80ºC.

#### Grp1 and nSH2

The gene encoding the Grp1 PH domain derived from human CYTH3 was expressed in BL21 (DE3) bacteria as a his6-MBP-N10-TEV-GGGG-Grp1-Cys fusion protein. The gene encoding the N-terminal SH2 (nSH2, 322-440aa) domain derived from the *PIK3R1* gene was cloned and expressed as a his6-GST-TEV-nSH2 fusion protein. A single cysteine was added to the C-terminus of the nSH2 domain to allow for chemical labeling with maleimide dyes. For both recombinant proteins, bacteria were grown at 37°C in 4 L of Terrific Broth for two hours or until OD_600_ = 0.8 and then shifted to 18°C for 1 hour. Cells were then induced to express either the Grp1 or nSH2 fusion by adding 0.1 mM IPTG. Cells were harvested 20 hours post-induction. We lysed bacteria into 50 mM Na_2_HPO_4_ [pH 8.0], 400 mM NaCl, 0.4 mM BME, 1 mM PMSF, 100 μg/mL DNase using a microfluidizer. Next, lysate was centrifuged at 16,000 rpm (35000 x g) for 60 minutes in a Beckman JA-20 rotor at 4ºC. Supernatant was circulated over 5 mL HiTrap chelating column (GE Healthcare, Cat# 17-0409-01) that was pre-incubated with 100 mM CoCl_2_ for 10 minutes, wash with MilliQ water, and equilibrated into lysis buffer lacking PMSF and DNase. Clarified cell lysate containing his6-MBP-N10-TEV-GGGG-Grp1-Cys was circulated over the HiTrap column and washed with 20 column volumes of 50 mM Na_2_HPO_4_ [pH 8.0], 300 mM NaCl, 0.4 mM BME containing buffer. Protein was eluted with buffer containing 50 mM Na_2_HPO_4_ [pH 8.0], 300 mM NaCl, and 500 mM imidazole at a flow rate of 4mL/min. Peak HiTrap elutant fractions were combined with 750 μL of 2 mg/mL TEV protease and dialyzed overnight against 4L of buffer containing 20 mM Tris [pH 8.0], 200 mM NaCl, and 0.4 mM BME. The next day, we recirculated cleaved proteins over two HiTrap (Co^+2^) columns (2 × 5 mL) that were equilibrated in 50 mM Na_2_HPO_4_ [pH 8.0], 300 mM NaCl, and 0.4 mM BME containing buffer for 1 hour. We concentrated the proteins via 10 kDa MWCO Vivaspin 20 to a volume of 5mL. The concentrated Grp1 protein was then loaded on a 124 mL Superdex 75 column equilibrated in in 20 mM Tris [pH 8], 200 mM NaCl, 10% glycerol, and 1 mM TCEP. Protein was eluted at a flow rate of 1mL/min. Peak fractions containing Grp1 were pooled and concentrated to 500-600 μM (∼8mg/mL). Peak fractions containing nSH2 were pooled and concentrated to 200-250 μM (∼3mg/mL). Proteins were frozen with liquid nitrogen and stored at -80ºC.

#### P-Rex1 (DH-PH) domain

The DH-PH domain of human P-Rex1 was expressed as a fusion protein, his6-MBP-N10-TEV-PRex1(40-405aa), in BL21(DE3) Star bacteria. Bacteria were grown at 37°C in 2L of Terrific Broth for two hours or until OD_600_ = 0.8. Cultures were shifted to 18°C for 1 hour then induced with 0.1 mM IPTG. Expression was allowed to continue for 20 hours before harvesting. Cells were lysed into buffer containing 50 mM NaHPO4 [pH 8.0], 400 mM NaCl, 5% glycerol, 1 mM PMSF, 0.4 mM BME, 100 μg/mL DNase using microtip sonication. Cell lysate was clarified by centrifugation at 16,000 rpm (35000 x g) for 60 minutes in a Beckman JA-20 rotor at 4ºC. To capture his_6_-tagged P-Rex1, cell lysate was circulated over a 5 mL HiTrap Chelating column (GE Healthcare, Cat# 17-0409-01) charged CoCl_2_. The column was washed with 100 mL of 50 mM NaHPO4 [pH 8.0], 400 mM NaCl, 5% glycerol, 0.4 mM BME buffer. Protein was eluted into 15 mL with buffer containing 50 mM NaHPO4 [pH 8.0], 400 mM NaCl, 500 mM imidazole, 5% glycerol, 0.4 mM BME. Peak fractions were pooled and combined with his6-TEV protease and dialyzed against 4 liters of buffer containing 50 mM NaHPO4 [pH 8.0], 400 mM NaCl, 5% glycerol, 0.4 mM BME. The next day, dialysate containing TEV protease cleaved protein was recirculated for 2 hours over a 5 mL HiTrap chelating column. Flowthrough containing P-Rex1 (40-405aa) was desalted into 20 mM Tris [pH 8.0], 50 mM NaCl, 1 mM DTT using a G25 Sephadex column. Note that some of the protein precipitated during the desalting step. Desalted protein was clarified using centrifugation followed by a 0.22μm syringe filter. P-Rex1(40-405aa, pI = 8.68) was further purified by cation exchange chromatography (i.e. MonoS) using a 20 mM Tris [pH 8.0], 0 – 1000 mM NaCl, 1 mM DTT. P-Rex1(40-405aa) bound eluted broadly in the presence of 100-260 mM NaCl. Pure fractions as determined by SDS-PAGE were pooled, concentration, and loaded onto a 120 mL Superdex 75 column equilibrated in 20 mM Tris [pH 8], 150 mM NaCl, 10% glycerol, 1 mM TCEP. Peak fractions containing P-Rex1(40-405aa) were pooled and concentrated to a concentration of 114 μM, aliquoted, frozen with liquid nitrogen, and stored at -80ºC.

#### Btk

The mutant Btk PI(3,4,5)P_3_ fluorescent biosensor was recombinantly expressed in BL21 Star *E. coli* as a his6-SUMO-Btk(1-171aa PH-TH domain; R49S/K52S)-SNAP fusion. Bacteria were grown at 37°C in Terrific Broth to an OD600 of 0.8. These cultures were then shifted to 18°C for 1 hr, induced with 0.1 mM IPTG, and allowed to express protein for 20 hr at 18°C before being harvested. Cells were lysed into 50 mM NaPO_4_ (pH 8.0), 400 mM NaCl, 0.5 mM BME, 10 mM Imidazole, and 5% glycerol. Lysate was then centrifuged at 16,000 rpm (35,172 × *g*) for 60 min in a Beckman JA-20 rotor chilled to 4°C. Lysate was circulated over 5 mL HiTrap Chelating column (GE Healthcare, Cat# 17-0409-01) charged with 100 mM CoCl_2_ for 2 hrs. Bound protein was then eluted with a linear gradient of imidazole (0–500 mM, 8 CV, 40 mL total, 2 mL/min flow rate). Peak fractions were pooled, combined with SUMO protease Ulp1 (50 μg/mL final concentration), and dialyzed against 4 L of buffer containing 20 mM Tris [pH 8.0], 200 mM NaCl, and 0.5 mM BME for 16–18 hr at 4°C. SUMO protease cleaved Btk was recirculated for 1 hr over a 5 mL HiTrap Chelating column. Flow-through containing Btk-SNAP was then concentrated in a 5 kDa MWCO Vivaspin 20 before being loaded on a Superdex 75 size-exclusion column equilibrated in 20 mM Tris [pH 8.0], 200 mM NaCl, 10% glycerol, 1 mM TCEP. Peak fractions containing Btk-SNAP were pooled and concentrated to a concentration of 30 μM before snap-freezing with liquid nitrogen and storage at –80°C. For labeling, Btk-SNAP was combined with a 1.5x molar excess of SNAP-Surface Alexa488 dye (NEB, Cat# S9129S) and incubated overnight at 4ºC. The next day, Btk-SNAP-AF488 was desalted into buffer containing 20 mM Tris [pH 8.0], 200 mM NaCl, 10% glycerol, 1 mM TCEP using a PD10 column. The protein was then spin concentrated using a Amicon filter and loaded onto a Superdex 75 column to isolate dye free monodispersed Btk-SNAP-AF488. The peak elution was pooled, concentrated, aliquoted, and flash frozen with liquid nitrogen.

#### p67/phox

Genes encoding the Rac1(GTP) biosensor, p67/phox, were cloned into a his10-TEV-SUMO plasmid and expressed in Rosetta2 (DE3) pLysS bacteria. We grew bacteria in 3L of Terrific Broth 37°C for two hours or until OD_600_ =0.8 before shifting temperature to 18°C for 1 hour. We induced protein expression in cells via addition of 50 μM IPTG. Cells expressed overnight for 20 hours at 18ºC before harvesting. We lysted cells into buffer containing 50 mM Na_2_HPO_4_ [pH 8.0], 400 mM NaCl, 0.4 mM BME, 1 mM PMSF, and 100 μg/mL DNase using a microfluidizer. The lysate was centrifuged at 16,000 rpm (35000 x g) for 60 minutes in a Beckman JA-20 rotor at 4ºC. Supernatant was then circulated over 5 mL HiTrap Chelating column (GE Healthcare, Cat# 17-0409-01) that was inoculated with 100mM CoCl_2_ for ten minutes. The HiTrap column was washed with 20 column volumes (100mL) of 50 mM Na_2_HPO_4_ [pH 8.0], 400 mM NaCl, 10 mM imidazole, and 0.4 mM BME containing buffer. Bound protein was eluted at a flow rate of 4mL/min with 15-20 mL of 50 mM Na_2_HPO_4_ [pH 8.0], 400 mM NaCl, and 500 mM imidazole containing buffer. Peak fractions were pooled and combined with his6-SenP2 (SUMO protease) at a final concentration of 50 μg/mL and dialyzed against 4 liters of buffer containing 25 mM Tris [pH 8.0], 400 mM NaCl, and 0.4 mM BME. Dialysate containing SUMO cleaved protein was recirculated for 2 hours over two 5 mL HiTrap Chelating (Co^2+^) columns that were equilibrated in buffer containing 25 mM Tris [pH 8.0], 400 mM NaCl, and 0.4 mM BME. Recirculated protein was concentrated to a volume of 5 mL using a 5 kDa MWCO Vivaspin 20 before loading on a 124 mL Superdex 75 column at a flow rate of 1mL/min. The column was equilibrated in buffer containing 20 mM HEPES [pH 7], 200 mM NaCl, 10% glycerol, and 1 mM TCEP. Peak fractions off the Superdex 75 column were concentrated in a 5 kDa MWCO Vivaspin 20 to a concentration between 200-500 μM (5-12mg/mL). Protein was frozen with liquid nitrogen and stored at -80ºC.

#### Farnesylated Gβ_1_/Gγ_2_ and SNAP-Gβ_1_/Gγ_2_

The native eukaryotic farnesyl Gβ_1_/Gγ_2_ and SNAP-Gβ_1_/Gγ_2_ complexes were expressed and purified from insect cells as previously described (Rathinaswamy et al. 2021; Kozasa and Gilman 1995; Hashem A. Dbouk et al. 2012). The Gβ_1_ and Gγ_2_ genes were cloned into dual expression vectors containing tandem polyhedron promoters. A single baculovirus expressing either Gβ_1_/his_6_-TEV-Gγ_2_ or SNAP-Gβ_1_/ his6-TEV-Gγ_2_ were used to infect 2-4 liters of High Five cells (2 x 10^6^ cells/mL) with 2% vol/vol of baculovirus. Cultures were then grown in shaker flasks (120 rpm) for 48 hours at 27ºC before harvesting cells by centrifugation. Insect cells pellets were stored as 10 g pellets in the -80ºC before purification. To isolate farnesylated Gβ_1_/his_6_-TEV-Gγ_2_ or SNAP-Gβ_1_/his6-TEV-Gγ_2_ complexes, insect cells were lysed by Dounce homogenization into 50 mM HEPES-NaOH [pH 8], 100 mM NaCl, 3 mM MgCl_2_, 0.1 mM EDTA, 10 μM GDP, 10 mM BME, Sigma PI tablets (Cat #P5726), 1 mM PMSF, DNase (GoldBio, Cat# D-303-1). We centrifuged homogenized lysate for 10 minutes at 800 x *g* to remove nuclei and large cell debris. We then centrifuged remaining lysate using a Beckman Ti45 rotor 100,000 x *g* for 30 minutes at 4 ºC. The post-centrifugation pellet containing plasma membranes was the resuspended in a buffer containing 50 mM HEPES-NaOH [pH 8], 50 mM NaCl, 3 mM MgCl2, 1% sodium deoxycholate (wt/vol, Sigma D6750), 10μM GDP (Sigma, cat# G7127), 10 mM BME, and a Sigma Protease Inhibitor tablet (Cat #P5726) to a concentration of 5 mg/mL total protein. We Dounce homogenized the sample to break apart membranes and then allowed the homogenized solution to stir for 1 hour at 4ºC. We centrifuged the solubilized extracted membrane solution in a Beckman Ti45 rotor 100,000 x *g* for 45 minutes at 4ºC. We diluted the supernatant containing solubilized Gβ_1_/his_6_-TEV-Gγ_2_ or SNAP-Gβ_1_/his6-TEV-Gγ_2_ in buffer composed of 20 mM HEPES-NaOH [pH 7.7], 100 mM NaCl, 0.1 % C_12_E_10_ (Polyoxyethylene (10) lauryl ether; Sigma, P9769), 25 mM imidazole, and 2 mM BME.

We affinity purified the soluble membrane extracted Gβ_1_/his_6_-TEV-Gγ_2_ or SNAP-Gβ_1_/his6-TEV-Gγ_2_ using Qiagen NiNTA resin. After adding NiNTA resin to the diluted solubilized extracted membrane solution, we allowed the resin to incubate and stir in a beaker at 4ºC for 2 hours. We packed our protein-bound resin beads into a gravity flow column and washed with 20 column volumes of buffer containing 20 mM HEPES-NaOH [pH 7.7], 100 mM NaCl, 0.1 % C_12_E_10_, 20 mM imidazole, and 2 mM BME. We eluted and discarded the G alpha subunit of the heterotrimeric G-protein complex by washing with warm buffer (30ºC) containing 20 mM HEPES-NaOH [pH 7.7], 100 mM NaCl, 0.1 % C_12_E_10_, 20 mM imidazole, 2 mM BME, 50 mM MgCl_2_, 10μM GDP, 30 μM AlCl_3_ (J.T. Baker 5-0660), and 10 mM NaF. We eluted Gβ_1_/his_6_-TEV-Gγ_2_ or SNAP-Gβ_1_/his6-TEV-Gγ_2_ form the NiNTA resin using buffer containing 20 mM Tris-HCl (pH 8.0), 25 mM NaCl, 0.1 % C_12_E_10_, 200 mM imidazole, and 2 mM BME. The eluted protein was incubated overnight at 4ºC with TEV protease to cleave off the his6 affinity tag.

The next day, the cleaved protein was desalted on a G25 Sephadex column into buffer containing 20 mM Tris-HCl [pH 8.0], 25mM NaCl, 8 mM CHAPS, and 2 mM TCEP. Next, we performed anion exchange chromatography using a MonoQ column equilibrated with the desalting column buffer. We eluted Gβ_1_/Gγ_2_ or SNAP-Gβ_1_/Gγ_2_ from the MonoQ column in the presence of 175-200 mM NaCl. Peak-containing fractions were combined and concentrated using a Millipore Amicon Ultra-4 (10 kDa MWCO) centrifuge filter. Concentrated samples of Gβ_1_/Gγ_2_ or SNAP-Gβ_1_/Gγ_2_, respectively, were loaded on either Superdex 75 or Superdex 200 gel filtration columns equilibrated 20 mM Tris [pH 8.0], 100 mM NaCl, 8 mM CHAPS, and 2 mM TCEP. Peak fractions were combined and concentrated in a Millipore Amicon Ultra-4 (10 kDa MWCO) centrifuge tube. Finally, we aliquoted the concentrated Gβ_1_/Gγ_2_ or SNAP-Gβ_1_/Gγ_2_ and flash frozen with liquid nitrogen before storing at -80ºC.

#### Fluorescent labeling of SNAP-Gβ_1_/Gγ_2_

To fluorescently label SNAP-Gβ_1_/Gγ_2_, protein was combined with 1.5x molar excess of SNAP-Surface Alexa488 dye (NEB, Cat# S9129S). SNAP dye labeling was performed in buffer containing 20 mM Tris [pH 8.0], 100 mM NaCl, 8 mM CHAPS, and 2 mM TCEP overnight at 4ºC. Labeled protein was then separated from free Alexa488-SNAP surface dye using a 10 kDa MWCO Amicon spin concentrator followed by size exclusion chromatography (Superdex 75 10/300 GL) in buffer containing 20 mM Tris [pH 8.0], 100 mM NaCl, 8 mM CHAPS, 1 mM TCEP. Peak SEC fractions containing Alexa488-SNAP-Gβ_1_/Gγ_2_ were pooled and centrifuged in a 10 kDa MWCO Amicon spin concentrator to reach a final concentration of 15-20 μM before snap freezing in liquid nitrogen and storing in the -80ºC. To calculate the SNAP dye labeling efficiency, we determined that Alexa488 contributes 11% of the peak A_494_ signal to the measured A_280_. Note that Alexa488 non-intuitively has a peak absorbance at 494 nm. We calculate the final concentration of Alexa488-SNAP-Gβ_1_/Gγ_2_ using an adjusted A_280_ (i.e. A_280(protein)_ = A_280(observed)_ – A_494(dye)_*0.11) and the following extinction coefficients: ε_280(SNAP-Gβ1/Gγ2)_ = 78380 M^−1^•cm^−1^, ε_494(Alexa488)_ = 71,000 M^-1^•cm^-1^.

### Fluorescent labeling of PI3K using Sfp transferase

As previously described (Rathinaswamy et al. 2021), we generated a Dyomics647-CoA derivative by incubating a mixture of 15 mM Dyomics647 maleimide (Dyomics, Cat #647P1-03) in DMSO with 10 mM CoA (Sigma, #C3019, MW = 785.33 g/mole) overnight at 23ºC. To quench excess unreacted Dyomics647 maleimide, we added 5 mM DTT. We thawed purified PIK3CB/ybbr-PIK3R1 (referred to as PI3Kβ or p110β-p85α in manuscript) and chemically labeled with Dyomics647-CoA using Sfp-his_6._ The ybbrR13 motif fused to PIK3R1 contained the following peptide sequence: D<u>S</u>LEFIASKLA (Yin et al. 2006). In a total reaction volume of 2mL we combined 5μM PI3Kβ, 4μM Sfp-his_6_, and 10μM DY647-CoA in buffer containing 20 mM Tris [pH 8], 150 mM NaCl, 10 mM MgCl_2_, 10% Glycerol, 1 mM TCEP, and 0.05% CHAPS. The ybbr labeling reaction was allowed to proceed for 4 hours on ice. Excess Dyomics647-CoA was removed via a using a gravity flow PD-10 column. We concentrated labeled Dy647-PI3Kβ in a 50 kDa MWCO Amicon centrifuge tube before loading on a Superdex 200 gel filtration column equilibrated in 20 mM Tris [pH 8], 150 mM NaCl, 10% glycerol, 1 mM TCEP, and 0.05% CHAPS (GoldBio, Cat# C-080-100). We pooled and concentrated peak fractions to 5-10μM before we aliquoted and flash froze with liquid nitrogen. Labeled protein was stored at -80ºC.

### Preparation of supported lipid bilayers

We generated small unilamellar vesicles (SUVs) for this study using the following lipids:: 1,2-dioleoyl-sn-glycero-3-phosphocholine (18:1 DOPC, Avanti # 850375C), 1,2-dioleoyl-*sn-*glycero-3-phospho-L-serine (18:1 DOPS, Avanti # 840035C), 1,2-dioleoyl-sn-glycero-3-phosphoethanolamine (18:1 (Δ9-Cis) DOPE, Avanti # 850725C), L-α-phosphatidylinositol-4,5-bisphosphate (Brain PI(4,5)P_2_, Avanti # 840046X), synthetic phosphatidylinositol 4,5-bisphosphate 18:0/20:4 (PI(4,5)P_2_, Echelon, P-4524), and 1,2-dioleoyl-sn-glycero-3-phosphoethanolamine-N-[4-(p-maleimidomethyl)cyclohexane-carboxamide] (18:1 MCC-PE, Avanti # 780201C). We report lipid mixtures as percentages equivalent to molar fractions. We dried a total of 2 μmoles lipids were combined with 2mL of chloroform in a 35 mL glass round bottom flask containing. This mixture was dried to a thin film via rotary evaporation where the glass round-bottom flask was kept in a 42ºC water bath. Following evaporation, we either flushed the lipid-containing flask with nitrogen gas or placed in it a vacuum desiccator for a minimum of 30 minutes. We obtained a concentration of 1 mM of lipids by resuspending the dried film in 2 mL of 1x PBS [pH 7.2]. We generated 30-50 nm SUVs from this 1 mM total lipid mixture via extrusion of the resuspended lipid mixture through 0.03 μm pore size 19 mm polycarbonate membrane (Avanti #610002) with filter supports (Avanti #610014) on both sides of the PC membrane. We prepared coverglass (25x75 mm, IBIDI, cat #10812) for depositing of SUV’s by first cleaning with heated (60-70ºC) 2% Hellmanex III (Fisher, Cat#14-385-864) in a glass coplin jar. We incubated hot Hellmanex III and coverglass for at least 30 minutes before rinsing with MilliQ water. The cleaned glass was then etched with Piranha solution (1:3, hydrogen peroxide:sulfuric acid) for 5-10 minutes. We rinsed and stored the etched coverglass in MilliQ. We rapidly dried our MilliQ-rinsed etched coverglass slides with nitrogen gas before adhering to a 6-well sticky-side chamber (IBIDI, Cat# 80608). We created SLBs by flowing 100-150μL of SUVs with a total lipid concentration of 0.25 mM in 1x PBS [pH 7.2] into the IBIDI chamber. Following 30 minutes of incubation, supported membranes were washed with 4 mL of 1x PBS [pH 7.2] to remove non-absorbed SUVs. To block membrane defects, we prepared 1 mg/mL beta casein (Thermo FisherSci, Cat# 37528) by clarifying with a centrifugation step at 4°C for 30 minutes at 21370 x *g* before passing through 0.22 μm syringe filtration unit (0.22 μm PES syringe filter (Foxx Life Sciences, Cat#381-2116-OEM). We then blocked membrane defects with 1 mg/mL beta casein (Thermo FisherSci, Cat# 37528) for 5-10 minutes.

When reconstituting amphiphilic molecules (i.e. lipids) in aqueous solution a variety of structures can form based on the lipid composition, including micelles, inverted micelles, and planar bilayers (Kulkarni 2019). The organization of these membrane structures is related to the molecular packing parameter of the individual phospholipids (Jacob N. Israelachvili, Mitchell, and Ninham 1976). The packing parameter 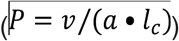 depends on the volume of the hydrocarbon 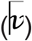, area of the lipid head group 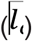, and the lipid tail length 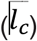. When generating supported lipid bilayers on a flat two-dimensional glass surface, we aim to create a fluid lamellar membrane. We find that phosphatidylcholine (PC) lipids are ideal for making supported lipid bilayers because they have a packing parameter of ∼1 (Costigan, Booth, and Templer 2000). In other words, PC lipids are cylindrical like a paper towel roll. In contrast, cholesterol and phosphatidylethanolamine (PE) lipids have packing parameters of 1.22 and 1.11, respectively (Angelov, Ollivon, and Angelova 1999; Carnie, Israelachvili, and Pailthorpe 1979). This gives cholesterol and PE lipids an inverted truncated cone shape, which prefers to adopt a non-lamellar phase structure under some condition. Due to the intrinsic negative curvature of PE lipids, they can spontaneously form inverted micelles (i.e. hexagonal II phase) in aqueous solution when they are the predominant lipid species (J. N. Israelachvili, Marcelja, and Horn 1980; Kobierski et al. 2022; Wnętrzak, Lątka, and Dynarowicz-Łątka 2013). In our hands, incorporation of PE lipids dramatically reduces the protein-maleimide coupling efficiency, displayed more membrane defects, and resulted in a larger fraction of surface immobilized Dy647-PI3Kβ. This could be related to the intrinsic negative curvature of PE membranes. However, further investigation is needed to decipher these issues.

### Protein conjugation of maleimide lipid

After blocking SLBs with beta casein, membranes were washed with 2 mL of 1x PBS and stored at room temperature for up to 2 hours before mounting on the TIRF microscope. Prior to single molecule imaging experiments, supported membranes were washed into TIRF imaging buffer. Supported membrane containing with MCC-PE lipids were used to covalently couple either Rac1(GDP) or phosphotyrosine peptide (pY). For the pY peptide experiments we used a doubly phosphorylated peptide derived from the mouse platelet derived growth factor receptor (PDGFR) with the following sequence: CSDGG(pY)MDMSKDESID(pY)VPMLDMKGDIKYADIE (33aa). The Alexa488-pY contained the same sequence with the dye conjugated to the C-terminus of the peptide. For these SLBs, 100 μL of 30 μM Rac1 was diluted in a 1x PBS [pH 7.2] and 0.1 mM TCEP buffer was added to the IBIDI chamber and incubated for 2 hours at 23ºC. Importantly, the addition of 0.1 mM TCEP significantly increases the coupling efficiency. SLBs with MCC-PE lipids were then washed with 2 mL of 1x PBS [pH 7.2] containing 5 mM beta-mercaptoethanol (BME) and incubated for 15 minutes to neutralize the unreacted maleimide headgroups. SLBs were washed with 1mL of 1x PBS, followed by 1 mL of kinase buffer before starting smTIRF-M experiments.

### Nucleotide exchange of Rac1

Membrane conjugated Rac1(GDP) was converted to Rac1(GTP) using either chemical activation (i.e. EDTA/ GTP/MgCl_2_) or the guanine nucleotide exchange factor (GEF), P-Rex1. Chemical activation was accomplished by washing supported membranes containing maleimide linked Rac1(GDP) with 1x PBS [pH 7.2] containing 1 mM EDTA and 1 mM GTP. Following a 15-minute incubation to exchange GDP for GTP, chambers were washed 1x PBS [pH 7.2] containing 1 mM MgCl_2_ and 50 μM GTP. A complementary approach that utilizes GEF-mediated activation of Rac1 was achieved by flowing 50 nM P-Rex1 DH-PH domain over Rac1(GDP) conjugated membranes (**Figure 1C**). Nucleotide exchange was carried out in buffer containing 1 × PBS, 1 mM MgCl_2_, 50 μM GTP. Both methods of activation yielded the same density of Rac1(GTP). Nucleotide exchange of membrane tethered Rac1 was assessed by visualizing the localization of the Cy3-p67/phox Rac1(GTP) sensor using TIRF-M.

### Single molecule TIRF microscopy

We preformed all supported membrane TIRF-M experiments in buffer containing 20 mM HEPES [pH 7.0], 150 mM NaCl, 1 mM ATP, 5 mM MgCl_2_, 0.5 mM EGTA, 20 mM glucose, 200 μg/mL beta casein (ThermoScientific, Cat# 37528), 20 mM BME, 320 μg/mL glucose oxidase (Biophoretics, Cat #B01357.02 *Aspergillus niger*), 50 μg/mL catalase (Sigma, #C40-100MG Bovine Liver), and 2 mM Trolox (Cayman Chemicals, Cat# 10011659). Perishable reagents (i.e. glucose oxidase, catalase, and Trolox) were added 5-10 minutes before starting image acquisition.

### Microscope hardware and imaging acquisition

Single molecule imaging experiments were performed at room temperature (23ºC) using an inverted Nikon Ti2 microscope using a 100x oil immersion Nikon TIRF objective (1.49 NA). We controlled the x-axis and y-axis position using a Nikon motorized stage, joystick, and Nikon’s NIS element software. We also controlled microscope hardware using Nikon NIS elements. Fluorescently labelled proteins were excited with one of three diode lasers: a 488 nm, a 561nm, or 637 nm (OBIS laser diode, Coherent Inc. Santa Clara, CA). The lasers were controlled with a Vortran laser launch and acousto-optic tuneable filters (AOTF) control. Excitation and emission light was transmitted through a multi-bandpass quad filter cube (C-TIRF ULTRA HI S/N QUAD 405/488/561/638; Semrock) containing a dichroic mirror. The laser power measured through the objective for single particle visualized was 1-3 mW. Fluorescence emission was captured on an iXion Life 897 EMCCD camera (Andor Technology Ltd., UK) after passing through one of the following 25 mm a Nikon Ti2 emission filters mounted in a Nikon emission filter wheel: ET525/50M, ET600/50M, and ET700/75M (Semrock).

### Kinetic measurements of PI(3,4,5)P_3_ lipid production

The phosphorylation of PI(3,4,5)P_3_ was measured on SLB’s formed in IBIDI chambers visualized via TIRF microscopy. We monitored the production of PI(3,4,5)P_3_ by solution-based PI3K at membrane surfaces using solution concentrations of 20 nM Btk-SNAP-AF488. Reaction buffer for experiments contained 20mM HEPES (pH 7.0), 150 mM NaCl, 5 mM MgCl_2_, 1 mM ATP, 0.1 mM GTP, 0.5 mM EGTA, 20 mM glucose, 200 μg/mL beta-casein (Thermo Scientific, Cat# 37528), 20 mM BME, 320 μg/mL glucose oxidase (Serva, #22780.01 *Aspergillus niger*), 50 μg/mL catalase (Sigma, #C40-100MG Bovine Liver), and 2 mM Trolox (Cayman Chemicals, Cat# 10011659). In experiments where inactive GTPases were coupled to membranes, no ATP was present in the reaction buffer and the 0.1 mM of GTP was replaced with 0.1 mM of GDP. Approximately 5-10 minutes before image acquisition, chemicals and enzymes needed the oxygen scavenging system were added to the TIRF imaging buffer.

The concentration of the Btk lipid sensor used for the kinetic assays does not interfere with the kinase activity. By comparing the membrane surface intensity of AF488-SNAP-Btk measured by TIRF microscopy in the presence of both 20 nM and near-saturating micromolar concentrations, we estimate that <0.1% of the PIP lipids are bound to a lipid sensor at any point during the kinetic experiments. Assuming an average footprint of 0.72 nm^2^ for phosphatidylcholine (Carnie, Israelachvili, and Pailthorpe 1979; Hansen et al. 2019), we calculated a density of 2.8 × 10^4^ PI(3,4,5)P_3_ lipids/μm^2^ for supported membranes that contain an initial concentrations of 2% PI(4,5)P_2_. We assume that the plateau fluorescence intensity of the AF488-SNAP-Btk sensor following reaction completion in the presence of PI3Kβ represents the production of 2% PI(3,4,5)P_3_. The bulk membrane intensity of AF488-SNAP-Btk was normalized from 0 to 1, and then multiplied times the total density of PI(3,4,5)P_3_ lipids to generate kinetic traces that report the kinetics of PI(3,4,5)P_3_ production.

We find that is critical to simultaneously visualize the localization of Btk-SNAP-AF488 and Dy647-PI3Kβ when performing lipid kinase assays. Poor quality supported membranes can artificially enhance PI3Kβ activity due to non-specific surface absorption of the kinase. When experiments displayed immobilized Dy647-PI3Kβ molecules, the data was discarded from our analysis and experiments were repeated.

### Surface density calibration

The density of membrane-tethered proteins attached to supported lipid bilayers was determined by coupling a defined ratio of either fluorescently labeled Cy3-Rac1 (1:10,000) or Alexa488-pY (1:30,000) in the presence of either 10 μM pY or 30 μM Rac1. The membrane surface density of GβGγ was quantified at equilibrium using a combination of AF488-SNAP-GβGγ (bulk signal) and dilute AF555-SNAP-GβGγ (0.0025%; 1:40,000), which allowed us to resolve and count the single molecule density. Single spatially resolved fluorescent proteins were visualize by TIRF microscopy. We calculated the single molecule densities of fluorescent labeled pY, Rac1, and GβGγ using ImageJ/Fiji Trackmake Plugin. The total surface density was calculated based on the dilution factor in the presence of dark unlabeled protein (e.g. Rac1(GTP), pY, or SNAP-GβGγ).

### Alphafold2 Multimer modelling

We utilized the AlphaFold2 using Mmseqs2 notebook of ColabFold at colab.research.google.com/github/sokrypton/ColabFold/blob/main/AlphaFold2.ipynb to make structural predictions of PI3Kβ (p110β/p85α) bound to G β γ. The pLDDT confidence values consistently scored above 90% for all models, with the predicted aligned error and pLDDT scores for all models are shown in **Figure 3 – figure Supplement 2**.

### Single particle tracking

Single fluorescent Dy647-PI3Kβ complexes bound to supported lipid bilayers were identified and tracked using the ImageJ/Fiji TrackMate plugin (Jaqaman et al. 2008). Data was loaded into ImageJ/Fiji as .nd2 files. We used the LoG detector to identify particles based on their size (∼6 pixel diameter), brightness, and signal-to-noise ratio. We then used the LAP tracker to generate trajectories that followed particle displacement as a function of time. Particle trajectories were then filtered based on Track Start (remove particles at start of movie), Track End (remove particles at end of movie), Duration (particles track ≥ 2 frames), Track displacement, and X - Y location (removed particles near the edge of the movie). The output files from TrackMate were then analyzed using Prism 9 graphing software to calculate the dwell times. To calculate the dwell times of membrane bound proteins we generated cumulative distribution frequency (CDF) plots with the bin size set to image acquisition frame interval (e.g. 52 ms). The log_10_(1-CDF) was plotted as a function dwell time and fit to a single or double exponential curve. For the double exponential curve fits, the alpha value is the percentage of the fast-dissociating molecules characterized by the time constant, τ _1_. A typical data set contained dwell times measured for n ≥ 1000 trajectories repeated as n = 3 technical replicates.Single exponential curve fit:

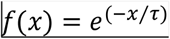

Two exponential curve fit:

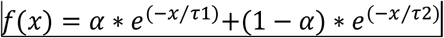

To calculate the diffusion coefficient (μm^2^/sec), we plotted probability density (i.e. frequency divided by bin size of 0.01 μm) versus step size (μm). The step size distribution was fit to the following models:

Single species model:

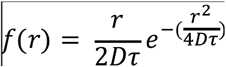

Two species model:

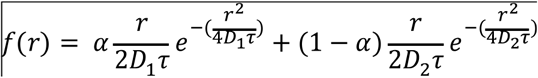

### Image processing, statistics, and data analysis

Image analysis was performed on ImageJ/Fiji and MatLab. Curve fitting was performed using Prism 9 GraphPad. The X-fold change in dwell time we report in the main text was calculated by comparing the mean single particle dwell time for different experimental conditions. Note that this is different from directly comparing the calculated dwell time (or exponential decay time constant, τ_1_). The X% reduction in diffusion or mobility we report in the main text was calculated by comparing the mean single particle displacement (or step size) measured under different experimental conditions.

