## Supplemental Figures for "Molecular dissection of PI3Kβ synergistic activation by receptor tyrosine kinases, GβGγ, and Rho-family GTPases"

### Figure 1 – figure supplement 1

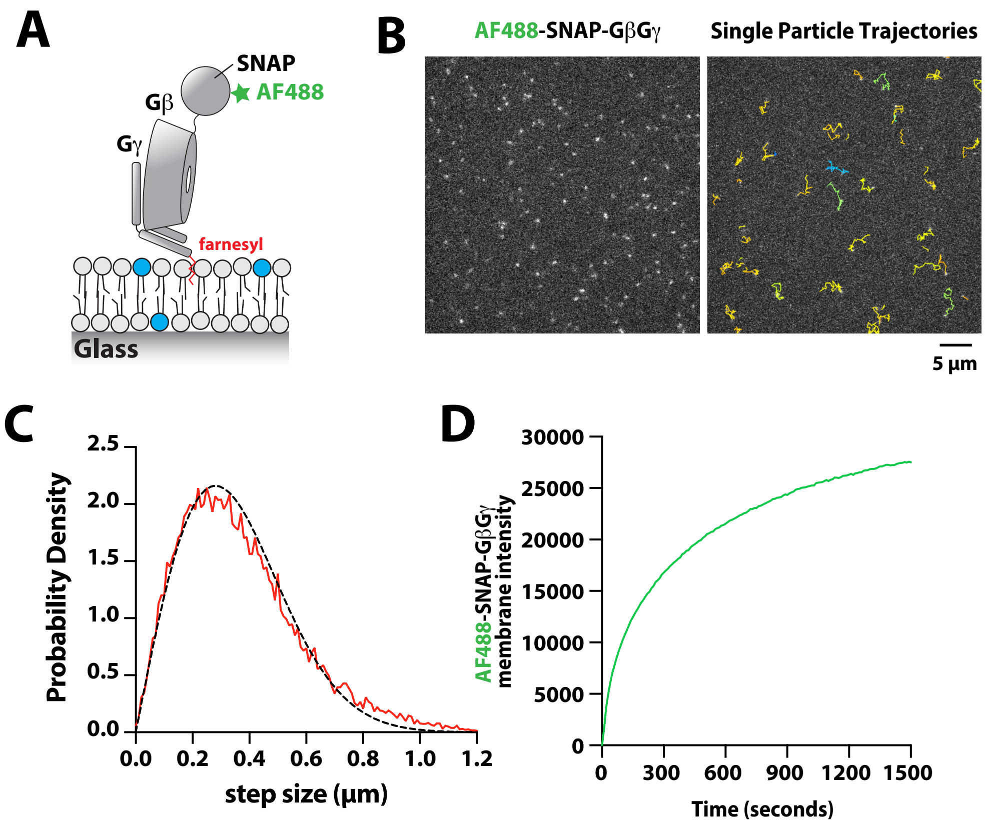


**Figure 1 – figure supplement 1**
**Characterization of Alexa488-SNAP-GβGγ localization on supported lipid bilayers**
(A) Cartoon showing the structural organization of the farnesylated GβGγ complex inserted into a supported lipid bilayer. The complex was labeled using Alexa488-SNAP surface dye, which we conjugated to the SNAP tag fused to the N-terminus of the Gβ subunit. (B) Representative TIRF-M images showing the localization of Alexa488-SNAP-GβGγ on supported lipid bilayer. The ImageJ TrackMate single molecule tracking plugin was used to identify and track Alexa488-SNAP-Gβ Gγ molecules. (C) Step size distribution showing single molecule displacements of N = 1128 Alexa488-SNAP-GβGγ particles (n = 23083 steps, D = 1.79 μm2/sec). (D) Bulk membrane absorption kinetics of 200 nM farnesyl GβGγ (10% Alexa488-SNAP-GβGγ). (C-D) Membrane composition: 98% DOPC, 2% PI(4,5)P2.

### Figure 1 – figure supplement 2

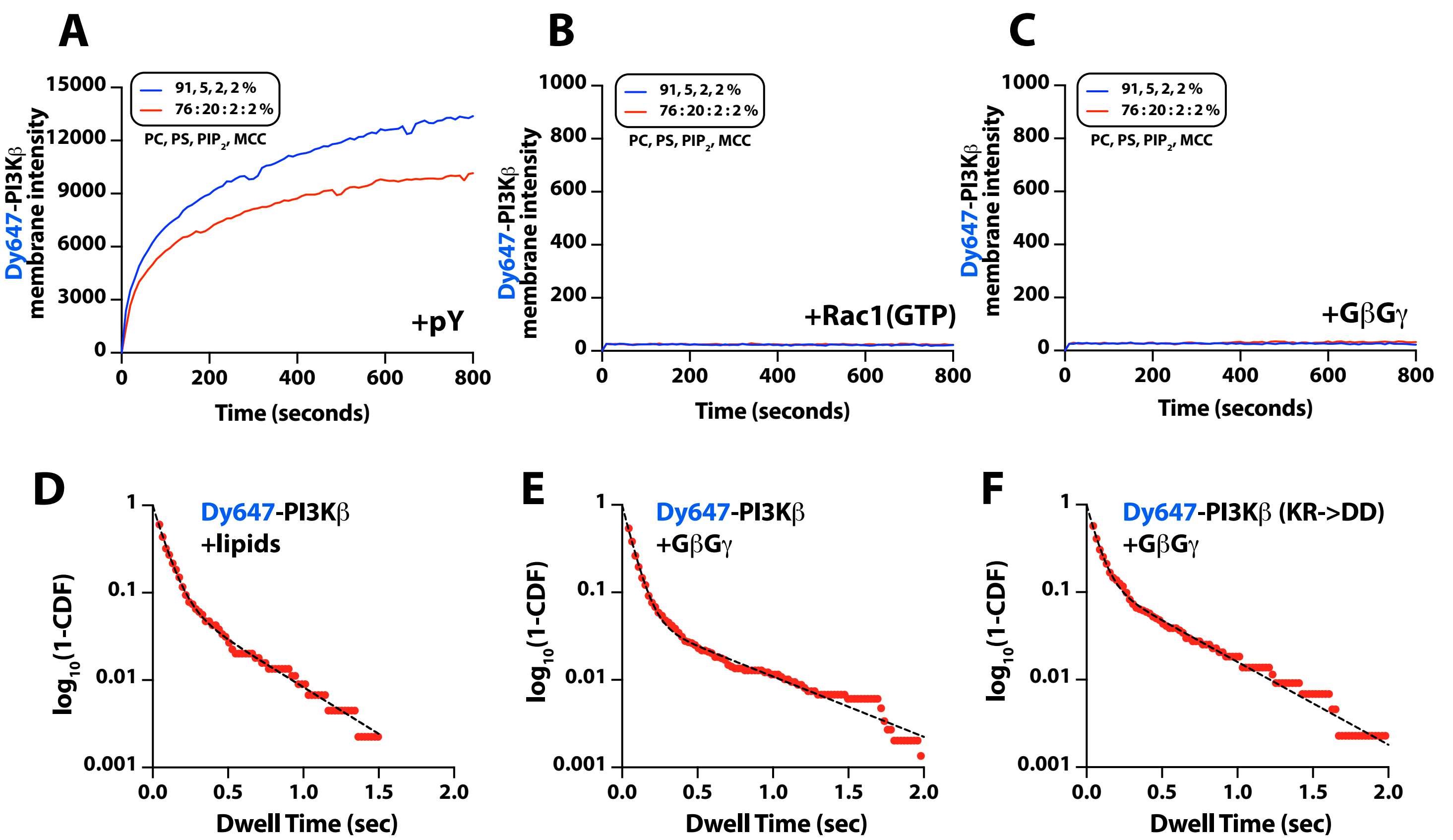


**Figure 1 – figure supplement 2**

**Characterization of Dy647-PI3Kβ membrane association with individual signaling inputs**

(A-C) Kinetic traces showing the bulk membrane localization of 10 nM Dy647-PI3Kβ measured of SLBs with either (A) pY, (B) Rac1(GTP), or (C) GβGγ tethered to the membrane. The following concentration of protein were used for membrane coupling or passive membrane absorption: (A) 10 μM pY, (B) 30 μM Rac1(GTP), or (C) 200 nM GβGγ. (A-C) Membrane lipid composition: 91% DOPC, 5% DOPS, 2% PI(4,5)P2, 2% MCC-PE (blue line); 76% DOPC, 20% DOPS, 2% PI(4,5)P2, 2% MCC-PE (red line). Note that the coupling efficiency of pY is generally reduced by high concentrations of anionic lipids, presumably due to electrostatic repulsion. (D-F) Single molecule dwell time distributions measured in the presence of (D) 10 nM Dy647-PI3Kβ + lipids (τ1=71±1ms, τ2=319±123ms, α=0.89, N=1821 particles) (E) 10 nM Dy647-PI3Kβ + 200 nM GβGγ (τ1=63±1ms, τ2=602±39ms, α=0.94, N=5992 particles), or (F) 10 nM Dy647-PI3Kβ GβGγ mutant (K532D/K533D) + 200 nM GβGγ (τ1=60±3ms, τ 2=494±49ms, α=0.87, N=2157 particles). Data plotted as log10(1–CDF) (cumulative distribution frequency). Dwell time statistics represent the average of n = 2 technical replicates. Alpha(α) represents the fraction of particles characterized by the time constant (τ1). Only particles tracked for at least two frames (44 ms) were include the distributions. Note that a 2000-fold higher concentration of Dy647-PI3Kβ is needed to observe single molecule binding events under these conditions compared to membrane containing membrane anchored pY peptide. (D-F) Membrane composition: 78% DOPC, 20% DOPS, 2% PI(4,5)P2.

### Figure 3 – figure supplement 1

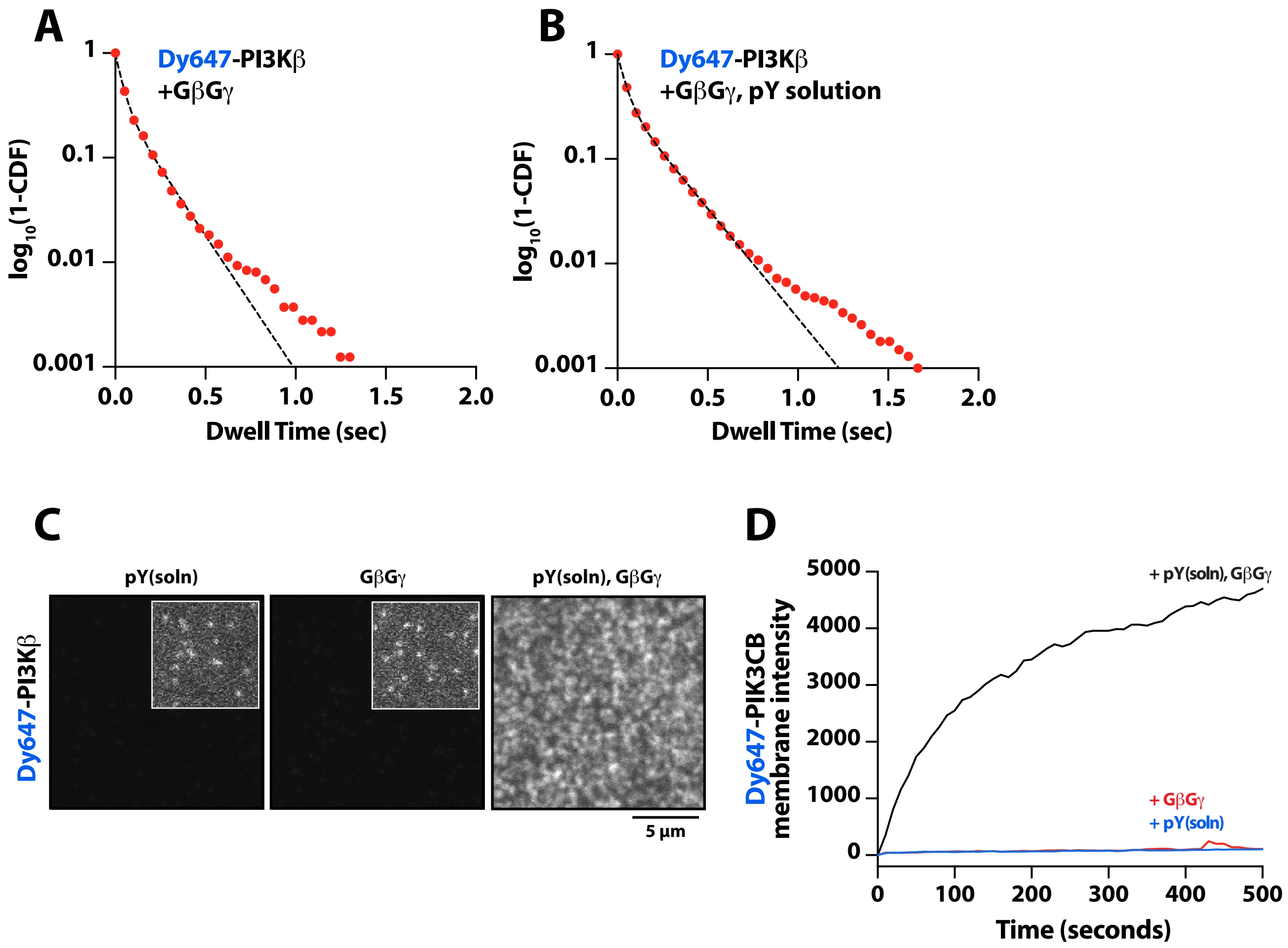


**Figure 3 – figure supplement 1**

**Solution pY peptide slightly enhances Dy647-PI3Kβ binding to GβGγ membranes**

(A-B) Single molecule dwell time distributions measured in the presence of 10 nM Dy647-PI3Kβ on supported membranes containing (A) GβGγ alone (τ1=40 ms, τ2=169 ms, α=0.65, N=3226 particles) or (B) GβGγ + 10 µM pY in solution (τ1=41 ms, τ2=193 ms, α=0.64, N=9639 particles). Data plotted as log10(1–CDF) (cumulative distribution frequency). Alpha(α) represents the fraction of particles characterized by the time constant (τ1). Note that a 2000-fold higher concentration of Dy647-PI3Kβ is needed to observe single molecule binding events under these conditions compared to membrane containing membrane anchored pY peptide. Membrane composition: 96% DOPC, 2% PI(4,5)P2, 2% MCC-PE. For these experiments, the MCC-PE lipids added for pY peptide conjugation was quenched with 5 mM beta-mercaptoethanol (BME). (C) Representative TIRF-M images showing the localization of 10 nM Dy647-PI3Kβ on supported lipid bilayers in the presence of either 20 µM pY (solution), 200 µM GβGγ (4,800 molecules/µm2), or pY(solution)/GβGγ. (D) Kinetic traces showing the bulk membrane localization of 10 nM Dy647-PI3Kβ measured of SLBs with either 20 µM pY (solution), 200 µM GβG γ (4,800 molecules/µm2), or pY(solution)/GβGγ. (C-D) Membrane composition: 78% DOPC, 20% DOPS, 2% PI(4,5)P2.

### Figure 3 – figure supplement 2

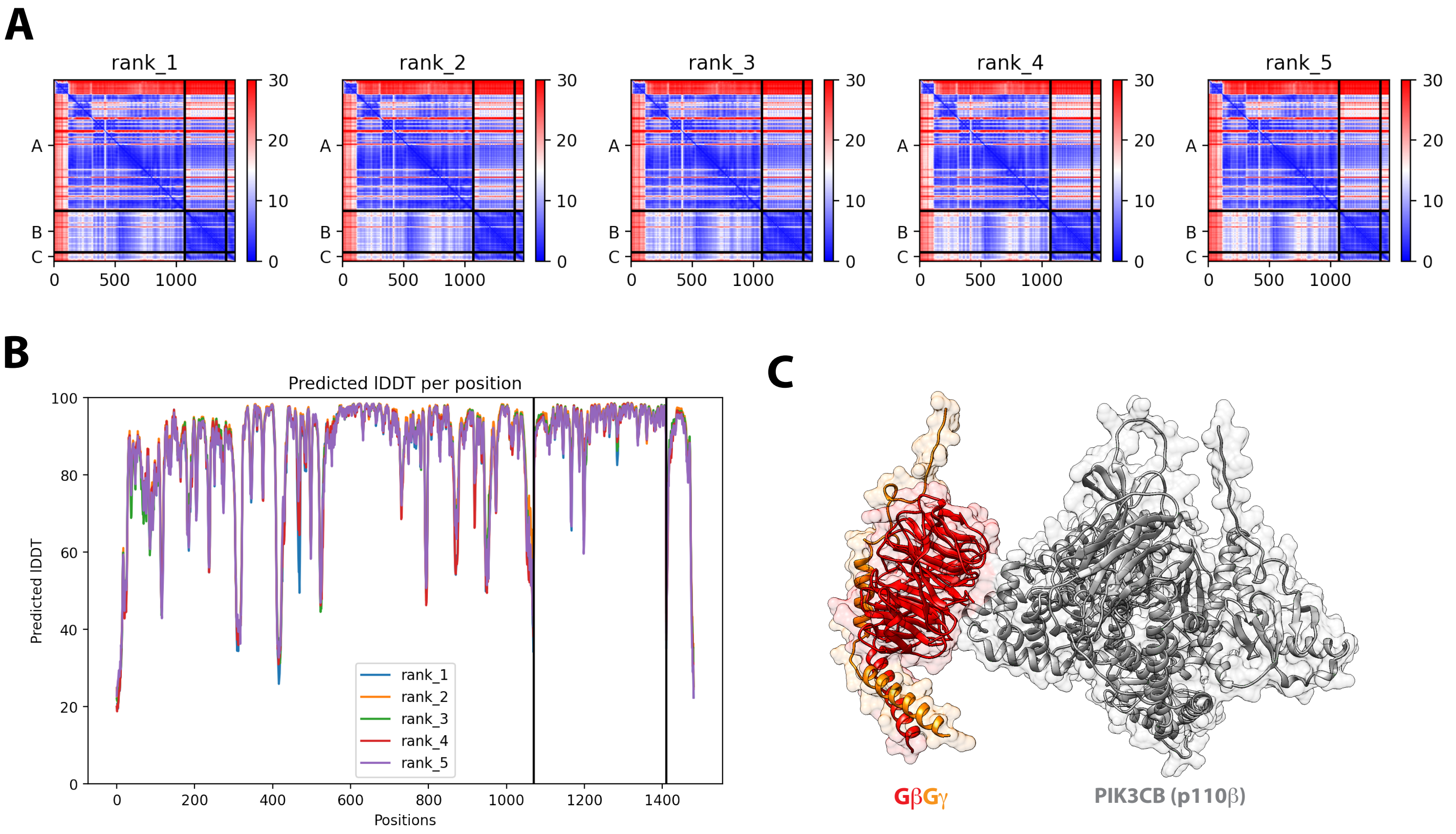


**Figure 3 – figure supplement 2**
**Structural model for GβGγ binding to the PI3Kβ catalytic subunit (p110β)**
(A) Predicted aligned error (PAE) for the top 5 ranked Alphafold2 Multimer structural prediction for full length p110β binding to GβGγ. (B) Predicted Local Distance Difference Test (IDDT) for the top 5 ranked Alphafold2 Multimer searches. (C) Alphafold2 Multimer structural model for GβGγ binding to p110β. The final model shown was generated in UCSF Chimera. All five models were consistent with experimental data from previous mutagenesis and HDX-MS data.

### Figure 5 – figure supplement 1

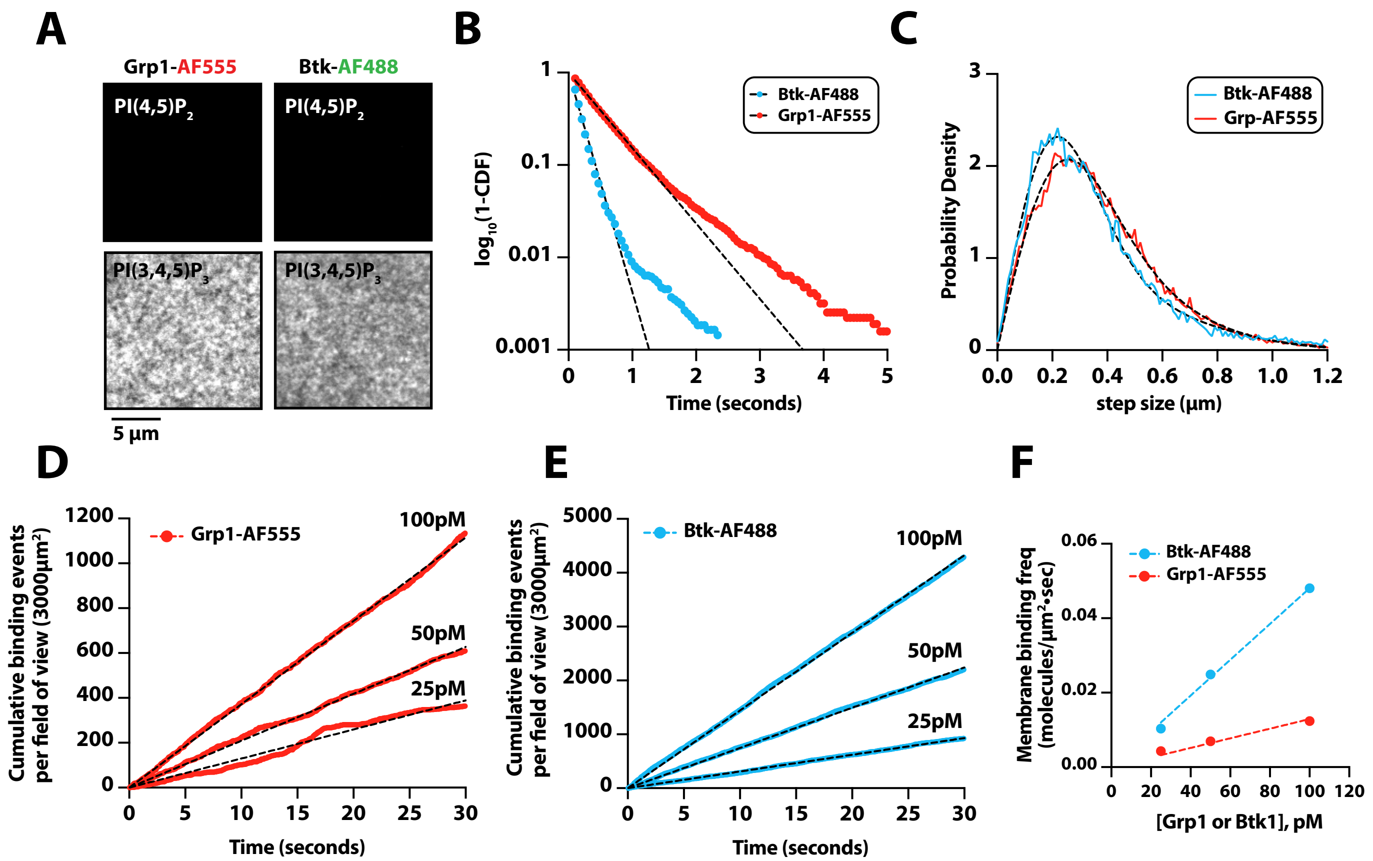


**Figure 5 – figure supplement 1**
**Characterization of Btk and Grp1 membrane binding**
(A) Representative TIRF-M images showing the localization of 50 nM Grp1-AF555 and 50 nM Btk-SNAP-AF488 on supported lipid bilayers containing either 2% PI(4,5)P2 or 2% PI(3,4,5)P3, plus 98% DOPC. (B) Single molecule dwell time distributions measured in the presence of 25 pM Btk-SNAP-AF488 (τ1= 183 ms, N=4865 particles) or 50 pM Grp1-AF555 (τ1= 523 ms, N=3160 particles). Dwell time distributions were fit to single exponential decay curves to calculate the characteristic dwell times (τ1). (C) Representative step-size distributions fit to a two species model for diffusion (dashed black line). Diffusivity was measured in the presence of either 25 pM Btk-SNAP-AF488 (D1 = 0.41 µm2/sec, D2 = 1.64 µm2/sec, α=0.66, N = 4865 particles, n = 25857 steps) or 50 pM Grp1-AF555 (D1 = 0.53 µm2/sec, D2 = 1.48 µm2/sec, α=0.6, N = 3160 particles, n = 34549 steps). (D-E) Cumulative membrane of binding events measured in the presence of 25 pM, 50 pM, and 100 pM (D) Grp1-AF555 or (E) Btk-SNAP-AF488. The field of view for the data collect in (D-E) equals 3000 µm2. (F) Calculation of the on rate (KON) for PI(3,4,5)P3 biosensors for data in (D-E). The slopes from each cumulative membrane binding curve in (D-E) were then plotted as a function of the Grp1-AF555 and Btk-SNAP-AF488 solution concentration to calculate the following association rate constants: KON(Grp1-AF555) = 0.13 Grp1 nM-1•µm-2•sec-1; KON(Btk-SNAP-AF488): 0.48 molecules nM-1•µm-2•sec-1. Membrane composition for single molecule measurements: 98% DOPC, 2% PI(3,4,5)P3.

### Figure 5 – figure supplement 2

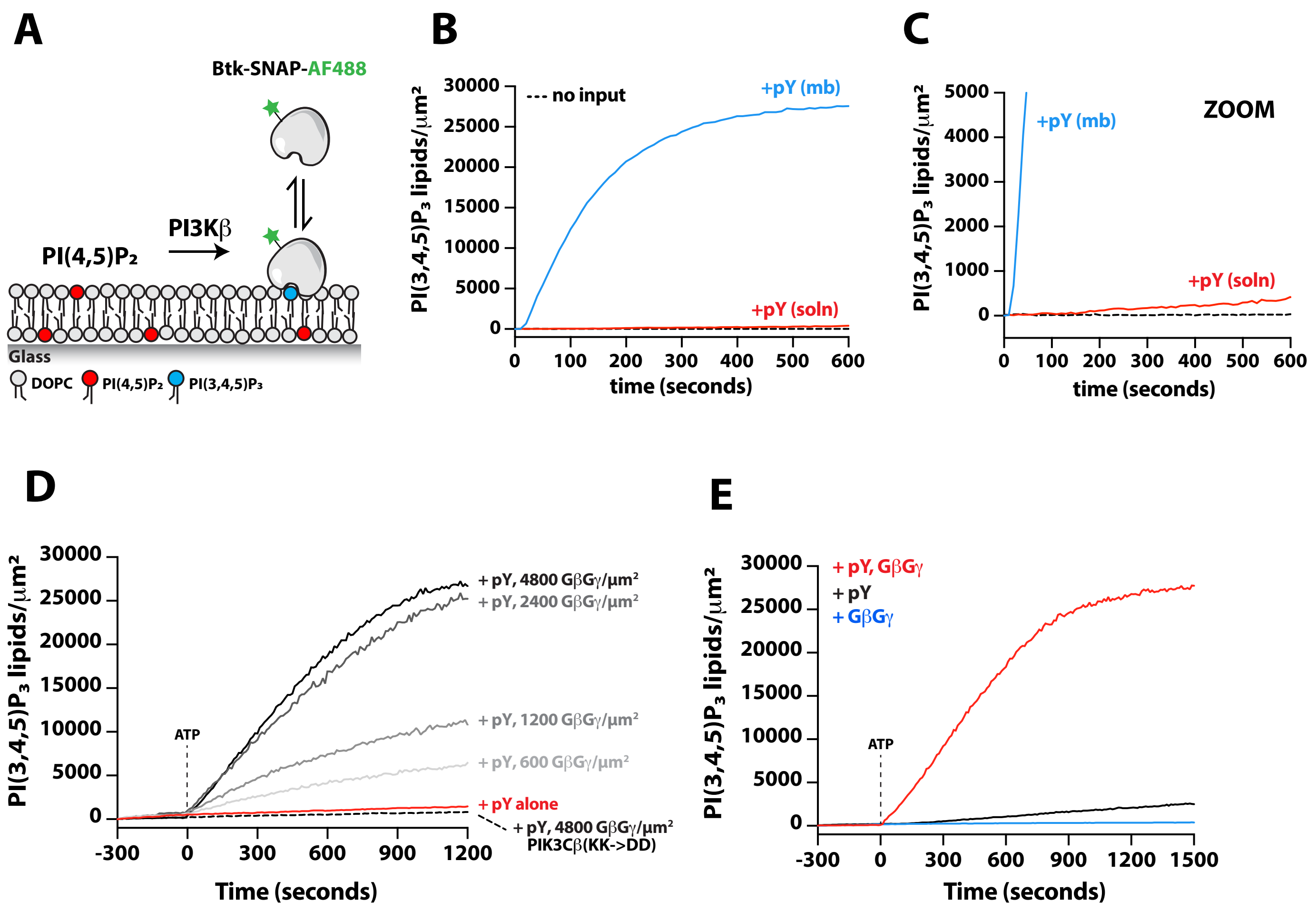


**Figure 5 – figure supplement 2**
**Reconstitution of PI3Kβ lipid kinase activity on supported membranes**
(A) Cartoon schematic showing how production of PI(3,4,5)P3 is monitored by visualizing membrane recruitment of soluble Btk-SNAP-AF488. (B) Kinetics of PI(3,4,5)P3 production monitored in the presence of 20nM Btk-SNAP-AF488 and 10 nM Dy647-PI3Kβ (note that this PI3Kβ concentration is higher than what was used to measure synergistic activation in Figure 5). Membranes were conjugated with 10 μM pY for 2 hours ("pY mb") or 10 μM pY was added in solution to stimulate Dy647-PI3Kβ. (C) Zoom in of graph in (B). In the presence of 10 nM Dy647-PI3Kβ, the initial reaction velocities were 157 PI(3,4,5)P3/μm2•sec (+pY mb) and 0.76 PI(3,4,5)P3/μm2•sec (+pY solution). Membrane composition: 96% DOPC, 2% PI(4,5)P2, 2% MCC-PE. (D) Kinetics of PI(3,4,5)P3 production monitored in the presence of 20nM Btk-SNAP-AF488, 20 pM Dy647-PI3Kβ, and the indicated densities of SNAP-GβGγ. Membranes contained either pY alone or pY/GβGγ. As a negative control, we measured the activity of the GβG binding mutant (Dy647-PI3Kβ K532D, K533D). Membrane composition: 96% DOPC, 2% PI(4,5)P2, 2% MCC-PE. (E) Kinetics of PI(3,4,5)P3 production monitored in the presence of 20nM Btk-SNAP-AF488 and 20 pM Dy647-PI3Kβ. Membrane contained either pY, Rac1(GTP), or GβGγ alone. Membrane composition: 76% DOPC, 20% DOPS, 2% PI(4,5)P2, 2% MCC-PE. (D-E) An ATP concentration of 1 mM was spiked into the reaction chamber at time zero to initiate the Dy647-PI3Kβ kinase activity measurements.
